# RANK-DEPENDENT CONTROL OF TUFT AND BEST4 CELL DEVELOPMENT IN THE INTESTINE

**DOI:** 10.1101/2025.08.18.670967

**Authors:** Reegan J. Willms, Tori McCabe, Lena Ocampo Jones, Ruth Schade, Edan A. Foley

## Abstract

Specialist intestinal epithelial cells are critical for barrier integrity and immune responses at the mucosal boundary, yet the pathways that govern their development are incompletely defined. Here, we identify an essential role for TNFRSF11A/RANK in shaping intestinal epithelial specialization in zebrafish. Using lineage trajectory analysis, we identified two tuft cell subtypes, including a subtype enriched for expression of genes required to produce pro-inflammatory leukotrienes. We showed that RANK deficiency reduced the abundance of immune-regulatory tuft and BEST4 cells, increased goblet cell frequency, and promoted the accumulation of pro-inflammatory leukocytes in the gut. Functionally, we demonstrated that BEST4 cell numbers expand following infection with a pandemic strain of *Vibrio cholerae*, and that RANK deficiency enhances fish susceptibility to host colonization by *Vibrio*, implicating this lineage in host defenses against an enteric pathogen. Together, our findings implicate RANK signaling in intestinal epithelial diversification and immune regulation.

## INTRODUCTION

Specialist intestinal epithelial cells control host immunity at the mucosal boundary. For example, crypt-resident Paneth cells release antimicrobial factors that contribute to niche sterility, while mucus-rich goblet cells maintain a physical barrier to microbial invasion, and M cells traffic luminal antigens to antigen-presenting cells in the lamina propria. More recent work suggests that the recently identified BEST4+ cell type may have important roles in regulation of luminal pH and mucosal barrier hydration(1,2), potentially adding to the rich mosaic of epithelial cells that protect from infection.

Within the community of immune-modulatory epithelial specialists, tuft cells are a relatively rare type that are primarily known for involvement in type two immunity(3). Upon detection of luminal helminths, tuft cells activate neighboring ILC2s, which then release IL-4 and IL-13 to promote hyperplastic generation of tuft and goblet cells that support the capture and peristaltic expulsion of helminths with the mucus barrier(4–6). Alongside established roles in type two immunity, recent studies uncovered involvement of tuft cells in antibacterial defenses(7), interactions with the commensal microbiome(8), and regeneration of damaged intestinal epithelia(9), indicating underexplored, yet foundational, involvements of tuft cells in gut function. Despite the importance of tuft cells for intestinal health, we know little about the signaling pathways that control tuft cell development and the relationships between tuft cells and their epithelial partners.

In this study, we investigated the role of TNFRSF11A/RANK in the generation of immune-regulatory epithelial cells in the zebrafish gut. RANK modifies patterning in several immunological structures(10), including the mammalian intestine, where RANK is essential for M cell development(11–13). However, RANK also controls gut remodeling during pregnancy(14); regulates ILC3 activity(15); and drives tuft cell expansion in mice challenged with *Nippostrongylus brasiliensis*(16), pointing to extensive involvements of RANK in maintenance of the epithelial barrier.

Zebrafish are an excellent system to characterize RANK-dependent control of intestinal epithelial development. The fish gut houses epithelial, mesenchymal, and hematopoietic cells that mirror those found in mammals(17,18), and there are striking similarities between the control of intestinal development in fish and mammals(19–22). We showed previously that RANK is expressed in larval intestinal progenitor cells, and that commensal bacteria induce *rank* expression in progenitors and BEST4 cells(23), indicating possible unexplored roles for RANK in fish intestinal epithelial development.

Here, we present evidence that RANK contributes to development of tuft and BEST4 cells in the zebrafish gut, and that whole-body loss of RANK leads to an accumulation of goblet cells, and induction of pro-inflammatory states in gut-associated leukocytes. Functionally, we establish that BEST4 cells significantly increase in number after a natural infection with diarrheagenic *Vibrio cholerae* and that loss of RANK enhances host colonization by *Vibrio*, suggesting potential roles for BEST4 cells in navigating enteric infections. Combined our work indicates that RANK contributes to development of the intestinal epithelial barrier and supports a role for BEST4 cells in host responses to pathogenic bacteria *in vivo*.

## RESULTS

### Zebrafish Intestines Contain Two Functionally Distinct Tuft Cell Types

In previous single cell gene expression atlases of zebrafish intestines(23,24), we identified a cluster of cells that expressed the definitive tuft cell marker *pou2f3* (Figure 1A-B). Upon re-examination, we discovered that this putative tuft cell population expressed multiple tuft cell markers such as *sh2d6*, tuft cell developmental regulators (*hmx2*, *sox9a*), type two immune receptors (*il4r.1* and *il13ra1*), and genes associated with leukotriene synthesis (*alox5a* and *alox5ap*) (Figure 1B). Like mammals, putative zebrafish tuft cells resolved into two distinct subtypes; one marked by expression of secretory gene products such as *arf1*, *sec22ba*, and *tmed2*, and a second subtype marked by expression of genes involved in immune responses such as STAT family transcription factors, *tnfrsf1a* and *nfkbiab* (Figure 1C-D). Of the two subtypes, the second was characterized by enriched expression of the IL13 receptor *il13ra1*, and genes required for production of pro-inflammatory leukotrienes, such as *alox5a*, *alox5ap* and *ltc4s* (Figure S1), suggesting that, like mammals, the zebrafish intestinal epithelium houses a tuft cell subtype that participates in type two immune defenses.

**Figure 1.**
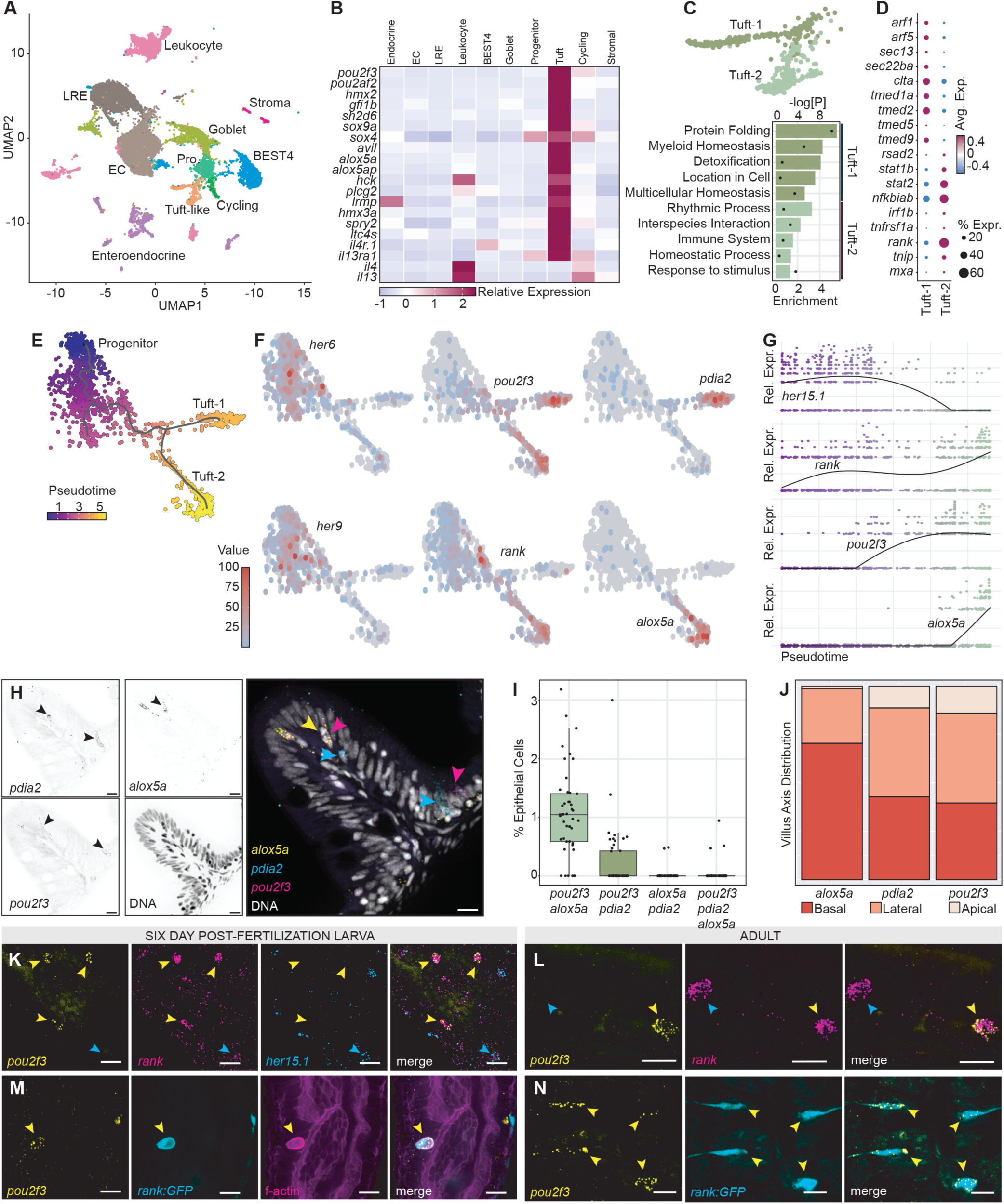
The Adult Zebrafish Has Two Tuft Cell Subtypes. **A:** UMAP representation of an adult single cell gene expression data set, indicating apricot-colored *pou2f3*-positive cells as “Tuft-like”. **B:** Heatmap representation of the relative expression of tuft-related genes in the indicated cell types. **C-D:** Sub-clustering resolves two tuft cell subsets that express genes related to the indicated Gene Ontology terms (C) and express the indicated markers (D) In (C) fold-enrichment is indicated by bar length and negative log p-values with circles. In (D), heatmap indicates scaled expression values. **E-G:** Visual representation of tuft cell maturation across pseudotime (E), including expression of the indicated marker genes (F-G). **H:** Fluorescence *in situ* hybridization showing the expression of the indicated genes in grayscale for each gene, and pseudo colored in the merged image. **I-J:** Percentage of intestinal epithelial cells that express the indicated markers (I), and position of the indicated cells along the villus axis (J). **K-N:** Visualizing expression of the indicated genes in wildtype (K-L), and Tg(*rank:GFP*, M-N) larval (K and M), and adult (L and N) zebrafish. In each image, scalebars are twenty micrometers, and arrowheads point to cells that express the gene of interest.

To better understand tuft cell development, we used lineage trajectory analysis to visualize tuft cell maturation over pseudotime (Figure 1E). Both tuft cell subtypes mapped back to a similar undifferentiated cell state characterized by Notch pathway activity, including expression of *her6*, *her9*, and the HES5 ortholog, *her15.1* (Figure 1E-G). As cells progressed along the trajectories and Notch activity declined, we observed an increase in *pou2f3* expression that persisted in both lineages, followed by subtype-specific expression of the Tuft-one marker *pdia2*, or the Tuft-two marker *alox5a* (Figure 1F-G).

To determine the abundance of the respective subtypes in the adult intestine, we performed fluorescence *in situ* hybridization assays for *pou2f3*, *pdia2*, and *alox5a* on sagittal intestinal sections of wildtype adult guts (Examples in Figure 1H and S2). We found that, like mammalian intestines, fish tuft cells are relatively rare, accounting for approximately 2% of all intestinal epithelial cells (Figure 1I). We observed minimal co-expression of *alox5a* and *pdia2*, suggesting that both genes mark distinct cell states (Figure 1I), and we found that *alox5a*- positive cells were slightly more prevalent at the villus base, while *pdia2*-positive cells appeared more evenly distributed along the villus axis (Figure 1J, Figure S2).

In addition to a putative *Notch*-*pou2f3*-*alox5a* trajectory, we were struck by early expression of the *tnfrsf11a*/*rank* gene during tuft cell development (Figure 1F-G). As *rank* modifies epithelial patterning in other tissues, we characterized the relationships between expression of *her15.1*, *rank*, and *pou2f3* in wildtype intestines, and in a reporter line that expresses GFP under control of 3.4kb of DNA upstream of the *rank* start site, Tg(*rank:GFP*)(23). In both instances, we confirmed overlapping expression of *rank* with *her15.1* and *pou2f3* (Figure 1K-L), as well as enriched expression of the *rank:GFP* reporter in *pou2f3*-positive epithelial cells (Figure 1M-N). Notably, *pou2f3*, *rank:GFP* double-positive cells were regularly characterized by actin-rich bundles at the cell periphery, a hallmark of mature tuft cells(25,26) (Figure 1M, arrowhead).

Thus, in combination with our prior TEM-based identification of intestinal epithelial cells with classical tuft cell morphology(23), our work shows that the zebrafish intestine has two rare tuft cell populations, including a specialist subtype that expresses type two immune receptors and genes required for production of pro-inflammatory leukotrienes.

### Comparisons Between Fish and Mouse Tuft Cells

In mice and humans, tuft cells arise from a crypt-associated progenitor that gradually transitions through a tuft-1 to a tuft-2 state as it migrates along the villus axis(27). As zebrafish intestines lack a crypt-like structure, and fish Tuft-1 and Tuft-2 subtypes are found throughout the epithelium, it is unclear to what extent the fish subtypes correlate with those found in mammals.

To characterize the relationship between tuft cells in fish and mammals, we compared the gene expression profiles of zebrafish tuft cells to recently described tuft-p, tuft-1 and tuft-2 populations from the mouse gut(27) (Figure 2A). We found that, like fish, *Rank* is expressed in mouse tuft cells, particularly in the progenitor population (Figure 2B), suggesting that *Rank* may have an evolutionarily conserved role in tuft cell development or function. Looking at the top eleven markers of fish Tuft-1 and Tuft-2 cell types, we discovered that eight markers from each population were also expressed in mouse tuft cells (Figure 2C-D), indicating broad transcriptional overlap between gene expression in tuft cells from both species. However, we also found that most fish Tuft-2 markers, particularly those linked to immune regulation and eicosanoid signaling (*Tm4sf4*, *Lgals2*, *Hopx*, *Alox5a*, *Alox5ap*) did not effectively distinguish between mouse tuft-1 and tuft-2 subtypes (Figure 2C). Instead, we found that zebrafish genes with restricted expression patterns among mouse tuft cells often marked the progenitor compartment, and are typically associated with the control of growth and inflammatory disease (e.g. *Myb*, *Krtcap2*, *Ckb*, Figure 2D). As a whole, these data indicate overlapping immune functions for fish and mouse tuft cells, but also suggest that fish subtypes may not directly correlate with mammalian counterparts.

**Figure 2.**
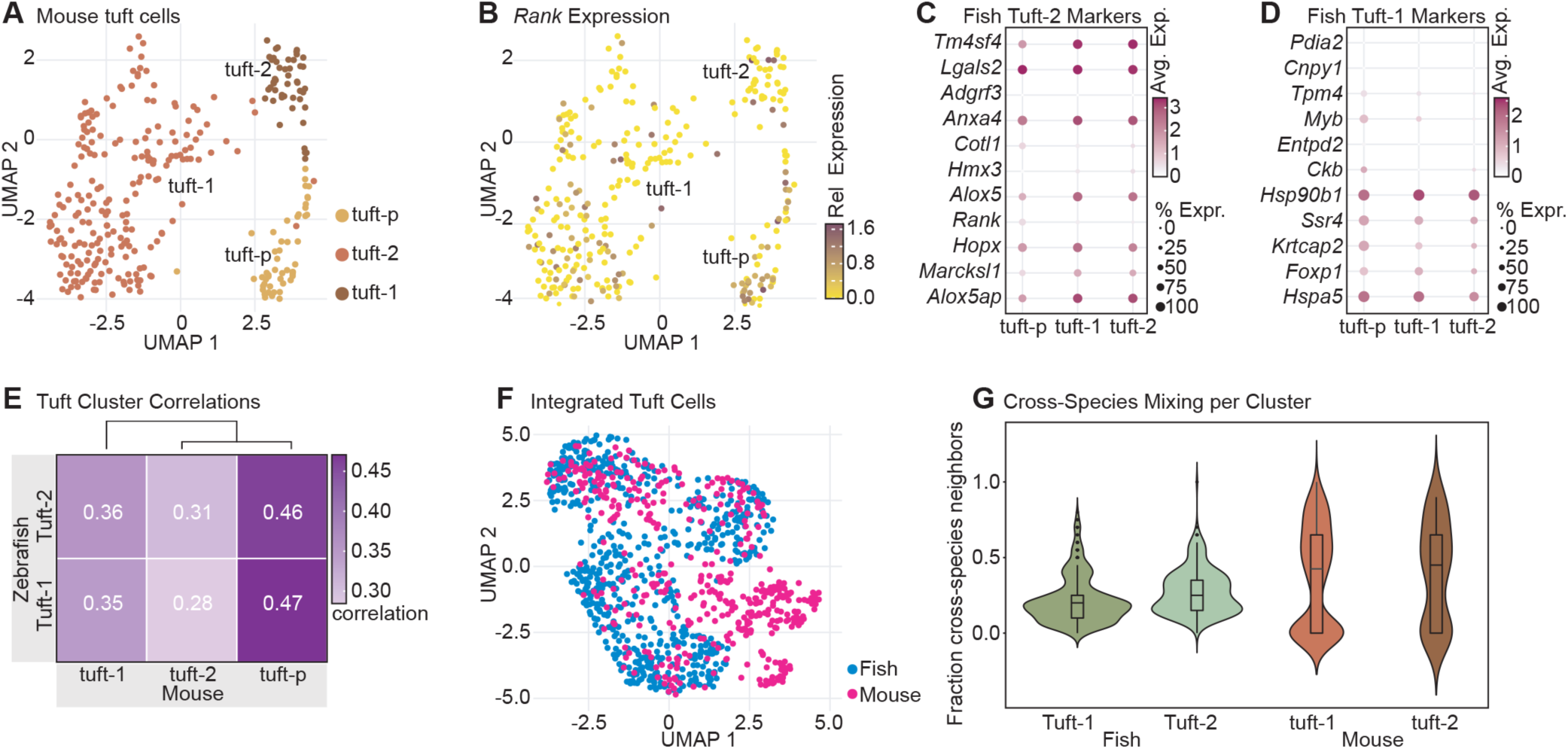
Transcriptomic comparison of fish and mouse tuft cell subtypes. **A-B:** UMAP representation of mouse tuft cells, indicating tuft-p, tuft-1 and tuft-2 subsets (A), alongside a feature plot showing expression of the *Rank* gene in mouse tuft cells (B). **C-D:** Dot plots showing the average expression of mouse orthologs of zebrafish Tuft-2 (C) and Tuft-1 (D) in the mouse tuft cell subsets. **E:** similarity matrix showing the degree of correlation between gene expression in zebrafish and mouse tuft cell subtypes. **F:** UMAP of cell clustering for mouse (fuchsia) and fish (deep sea blue) tuft cells. **G:** Violin plot showing the fraction of cross-species neighbors for fish and mouse tuft cells in panel F.

To better assess relatedness between tuft cells in the two animals, we then determined the extent to which the gene expression profiles of fish tuft cell subtypes correlate with those of mouse tuft cell subtypes. These analyses revealed that both zebrafish tuft cell subtypes correlate most closely with the mouse tuft-p population (Figure 2E), indicating that zebrafish tuft cells more closely resemble a mammalian progenitor-like tuft gene expression program than specialized tuft-1 and tuft-2 cell types. Through integration of the fish and mouse tuft cell gene expression data sets (Figure 2F) and quantification of cross-species neighbor mixing (Figure 2G) we further confirmed overlap between the two animals, but also observed species separation, adding further weight to the hypothesis that fish and mouse tuft cells have shared transcriptional programs, but also animal-specific components. Combined, these data suggest that fish and mouse tuft cells have evolutionarily conserved immune-regulatory functions, but also indicate significant differences in gene expression patterns between the two animals.

### Rank Promotes Tuft Cell Development

As *rank* expression marks a putative early tuft cell population in fish and mice (Figure 1E, Figure 2B), we next elected to characterize RANK function in greater detail. Transcriptionally, we observed pronounced *rank* expression in epithelial progenitor (Figure 3A, 36% of all progenitors) and tuft cell populations (44%), with lesser expression in goblet cells (7%) and BEST4 cells (6%), as well as minimal expression in Enterocytes (1%). These data support possible roles for RANK in tuft cell function or development and match our observation that commensal microbes promote *rank* expression in larval progenitor and BEST4 cells(23).

**Figure 3.**
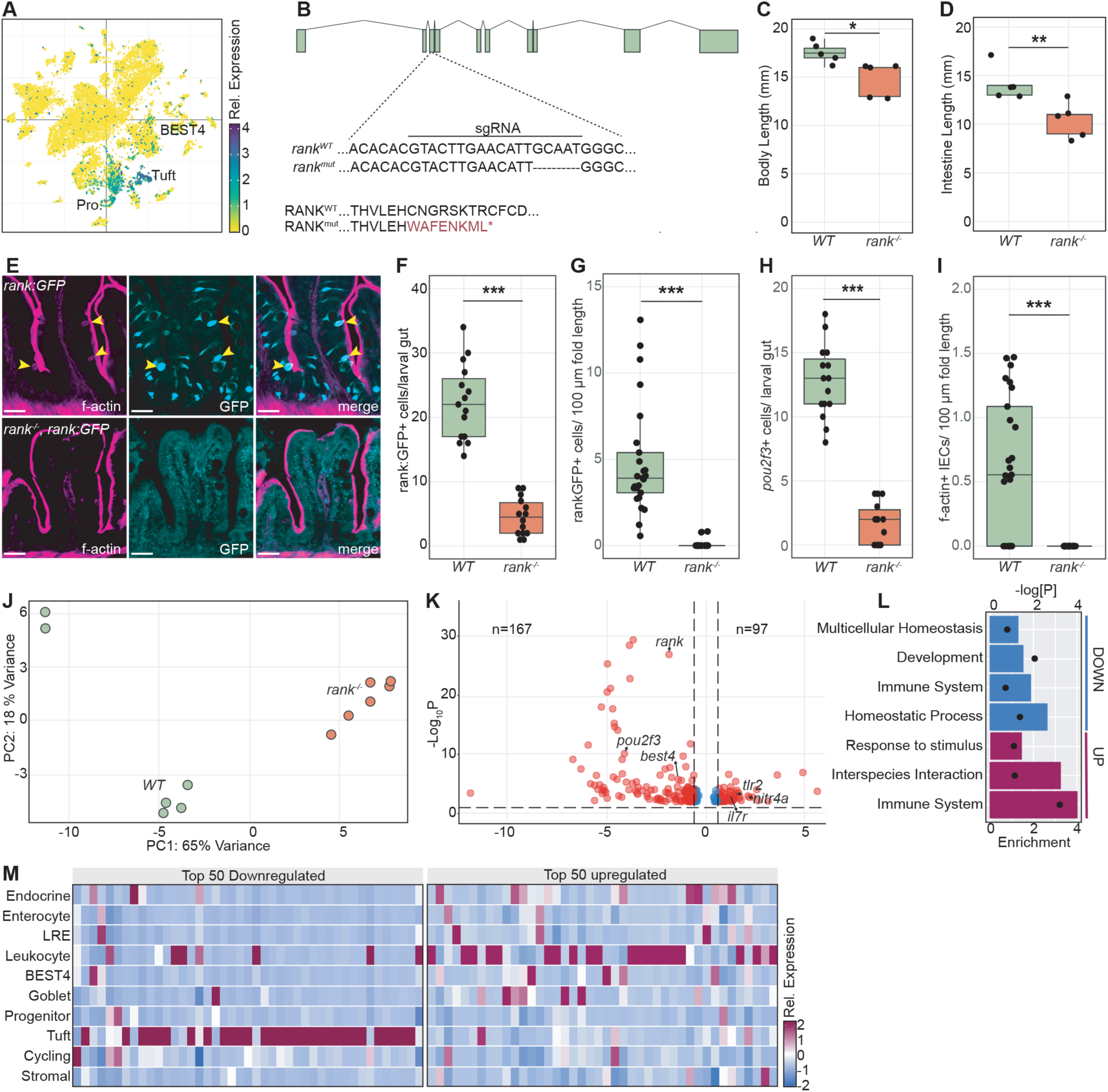
RANK Deficiency Impacts Intestinal Immunity. **A:** Feature plot showing *rank* gene expression levels in adult zebrafish intestines. **B:** Schematic representation of the *rank* gene locus, with exonic DNA indicated in sage. The genomic region targeted by a RANK-directed sgRNA is indicated below, and the wildtype sequence alongside the mutated sequence is presented at the bottom. **C-D:** Body length (B), and intestinal (C) length of wildtype and RANK-deficient siblings. **E-I:** Visualization of filamentous actin (magenta) and GFP (cyan) in wildtype or *rank* mutant *rank:GFP* fish. The numbers of GFP+ cells per larval (F) and adult (G) guts are shown, as well as the number of *pou2f3*-positive larval (H) and f-actin rich (I) adult cells. **J-L:** Principal Component Analysis plot (J) of gene expression in wildtype and *rank* siblings. Horizontal dashed line in the volcano plot (K) shows significantly differentially expressed (padj < 0.01) genes in mutants relative to wildtype fish, and the corresponding down and up-regulated gene ontology terms are shown in (L). Enrichment is indicated by bar length and negative log p-values with circles. **M:** Heatmap representation of the relative expression of the top 50 up and down-regulated genes in the indicated cell types. In each panel one asterisk = p<0.05, two = P < 0.01, and three = P < 0.001, calculated by ANOVA. In each image, scalebars are twenty micrometers, and arrowheads point to cells that express the gene of interest.

To assess the role of RANK in intestinal homeostasis, we used CRISPR mutagenesis to engineer a fish line with a five-nucleotide deletion in the third exon, generating a truncated protein with a frameshift at H58, followed by eight out of frame amino acids, and a premature stop codon (Figure 3B). The resulting protein lacks the C-terminal portion of the ligand binding domain, alongside the entire transmembrane and intercellular domains. Although homozygous viable, *rank* mutants were smaller than wildtype controls (Figure 3C), with significantly shorter intestines (Figure 3D), and impaired intestinal NF-kB activity (Figure S3). Homozygous *rank* mutants showed progressive symptoms of scoliosis with age (Figure S4), consistent with an essential role for RANK in osteoclast function(28). We found that *rank* deficiency abolished GFP expression in *rank:GFP* fish (Figure 3E-G), led to a significant loss of *pou2f3*-positive cells in the larval gut (Figure 3H), and an absence of actin-rich cells throughout the intestinal epithelium (Figure 3E, I), suggesting that RANK deficiency impacts the generation of mature tuft cells.

To better assess the impacts of RANK deficiency on gut function, we performed bulk RNA sequencing analysis of gene expression in intestines dissected from eight-week-old, co-housed, wildtype and *rank* mutant siblings derived from a heterozygous in-cross. Principal component analysis showed that *rank* mutant intestines were clearly distinct from wildtype counterparts (Figure 3J), with significant loss of expression of *rank* and *pou2f3* (Figure 3K), as well as disrupted expression of multiple genes linked with immune regulation (Figure 3L). To identify the intestinal cell types that are most sensitive to loss of *rank*, we mapped the fifty most significantly up and down-regulated genes in *rank* mutants to our adult single cell gene expression atlas(24). In this way, we discovered that greater than 70% of the most downregulated genes marked tuft cells, while 50% of the most upregulated genes marked leukocytes (Figure 3M), suggesting impaired immune activity in RANK-deficient guts hallmarked by diminished tuft cell function, and enhanced hematopoietic cell activity.

### RANK Deficiency Impacts the Functional State of Gut-Associated Leukocytes

As RANK modifies leukocyte-epithelium interactions in mice (15,29), and loss of RANK deregulated the expression of leukocyte-associated immune effectors in the zebrafish gut (Figure 3M), we next characterized the consequences of RANK deficiency for the intestinal hematopoietic compartment. Here, we used our adult single cell gene expression atlas(24), to resolve the wildtype expression patterns of all leukocyte-associated genes that were dysregulated in *rank* mutants. Consistent with Figure 3M, 90% of leukocyte-associated genes were expressed to higher levels in *rank* mutants than in their wildtype siblings (Figure 4A). Of the few downregulated genes, we observed significantly diminished expression of the ILC2 marker *il13*, alongside lower expression of several dendritic cell-associated genes (Figure 4B), matching previous results that signals from tuft cells prompt the activation of gut-associated ILC2s and dendritic cells(5,30). In contrast, we noted broadly enhanced expression of genes essential for antigen presentation (e.g. *cd74a* and *cd74b*), as well as increased expression of immune effectors specific to ILC3s (e.g. *nitr5* and *nitr4a*), macrophages (e.g. *mpeg1.1* and *cxcl19*), B cells (e.g. *tlr3* and *flt3*), and T cells (e.g. *cd4-1* and *il7r*) (Figure 4B), indicating enhanced immune activity within gut-associated myeloid and lymphoid lineages in *rank* mutant fish.

**Figure 4.**
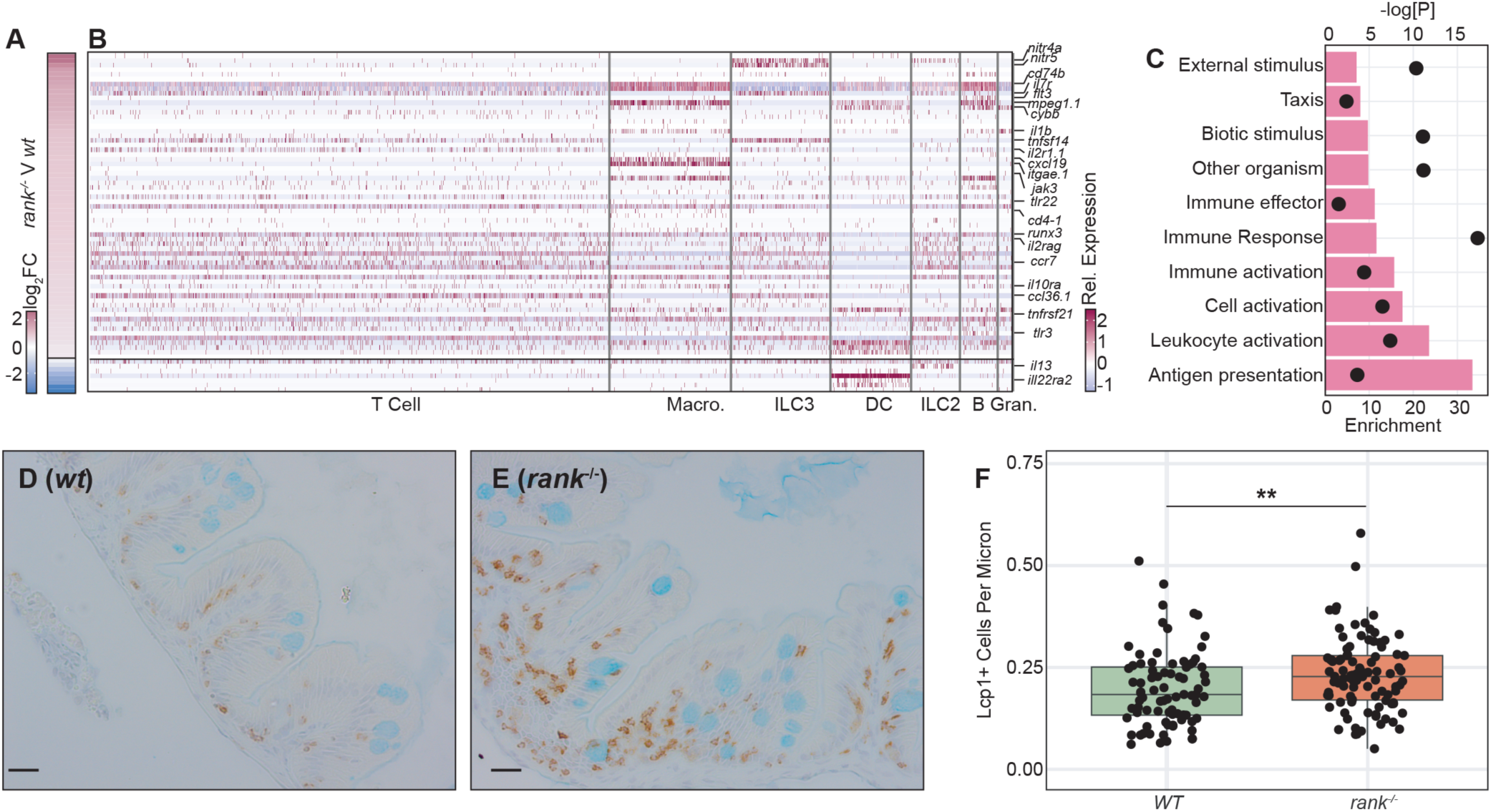
RANK Deficiency Enhances Intestinal Infiltration by Activated Leukocytes: **A-B:** Expression of all leukocyte-associated genes in *rank* mutants relative to wildtype siblings (A), and cell-type expression patterns of each gene. **C:** Gene ontology analysis of functions associated with all leukocyte-associated genes with enriched expression in *rank* mutants. Fold-enrichment is indicated by bar length and negative log p-values with circles **D-F:** Visualization (D-E) and quantification (F) of LCP-1-positive leukocytes (burnt sienna) in the intestines of wildtype (wt) and *rank* mutant siblings that were counterstained with Alcian blue to visualize goblet cells (aquamarine). Two asterisks = P < 0.01 calculated by ANOVA. Scalebars are twenty micrometers.

GO term analysis of all leukocyte-associated genes that are dysregulated in *rank* mutants confirmed that RANK deficiency switched leukocytes to an active, pro-inflammatory state, marked by elevated expression of genes required for leukocyte activation, antigen presentation, and immune effector functions (Figure 4C). To directly examine effects of RANK deficiency on gut-associated leukocytes, we then stained sagittal sections of intestines prepared from co-housed eight-week-old wildtype and RANK-deficient siblings for the pan-leukocyte marker LCP-1 (Figure 4D-E). In this experiment, we detected a mild, but significantly enhanced recruitment of LCP-1-positive cells in *rank* mutant intestines relative to wildtype controls (Figure 4F), and we found that leukocytes from RANK-deficient fish consistently had enhanced levels of LCP-1 staining (Figure 4D-E). In sum, our work shows that loss of RANK results in a moderate accumulation of intestinal leukocytes that are characterized by enhanced expression of activation markers, indicating that RANK-deficiency shifts the hematopoietic compartment to a pro-inflammatory state in adult zebrafish.

### RANK Promotes Development of *alox5a*-positive Tuft Cells

As RANK deficiency also impacted expression of tuft cell markers, we next used fluorescence *in situ* hybridization to visualize expression of *rank*, *alox5a*, and *pdia2* in anterior, middle, and posterior intestinal segments(31) of eight-week-old wildtype (Figure 5A) and *rank* mutant siblings (Figure 5B). Consistent with our transcriptomic data, we did not detect *rank* transcript in *rank* mutant intestines (Figures 5A-B). In contrast, we observed regionalized impacts of RANK deficiency on the development of mature tuft cell subtypes. For example, whereas loss of *rank* led to a moderate decline in *pdia2*-positive tuft cells exclusively in the intestinal anterior region, we discovered that RANK deficiency led to significant loss of *alox5a*-positive type-two tuft cells throughout all three intestinal epithelial segments (Figure 5C-E), indicating an intestine-wide role for RANK in the development of *alox5a*+ immune-regulatory tuft cells in the adult intestine, and a regionalized role for RANK in the specification of *pdia2+* tuft cells in the anterior intestinal segment.

**Figure 5.**
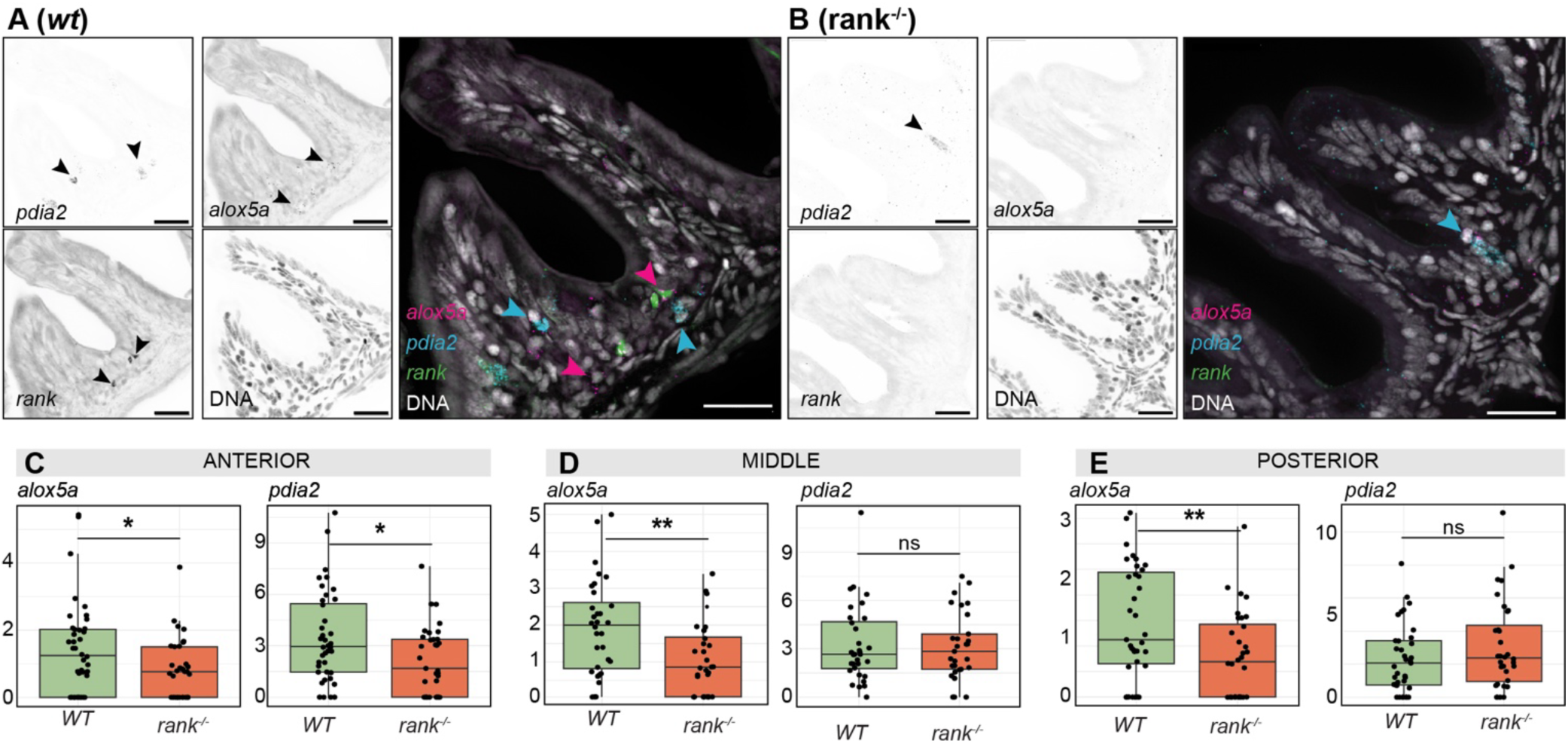
RANK Deficiency Results in Intestinal Loss of Type-2 Tuft Cells. **A-B:** Fluorescence *in situ* hybridization images of sagittal intestinal sections from wildtype (wt) and *rank* mutant siblings showing the expression of the indicated genes in grayscale for each gene, and pseudo colored in the merged image. **C-D:** Quantification of *alox5a* and *pdia2*-positive cells in anterior middle and posterior intestinal sections of wildtype (wt) and *rank* mutant siblings. In each panel one asterisk = p<0.05, and two = P < 0.01, calculated by ANOVA. In each image, scalebars are twenty micrometers, and arrowheads point to cells that express the gene of interest.

### Loss of RANK Diminishes Expression of BEST4 Cell Markers

Thus far, our work suggests that loss of RANK impacts the functional states and relative abundances of two established modifiers of intestinal immunity: tuft cells and gut-associated leukocytes. However, we were also surprised by an unexpected decline in the expression of multiple BEST4 cell markers in *rank* mutants, including significantly reduced expression of definitive BEST4 cell markers such as *cftr*, *otop2*, and *best4* (Figure 6A). BEST4 cells have only been recently described but are common in the animal kingdom with the somewhat surprising exception of mice(32), and we and others previously showed that larval and adult zebrafish intestines include related populations of BEST4 cells that express classical BEST4 cell markers(23,24,33,34), including *guanylate cyclase 2C*, ion channels, and carbonic anhydrases (Figure 6B), Unlike mammals, zebrafish BEST4 cells do not appear to express *spi2* or *spic*(35), the expression of which marks dendritic cells in the adult zebrafish gut (Figure S5).

**Figure 6.**
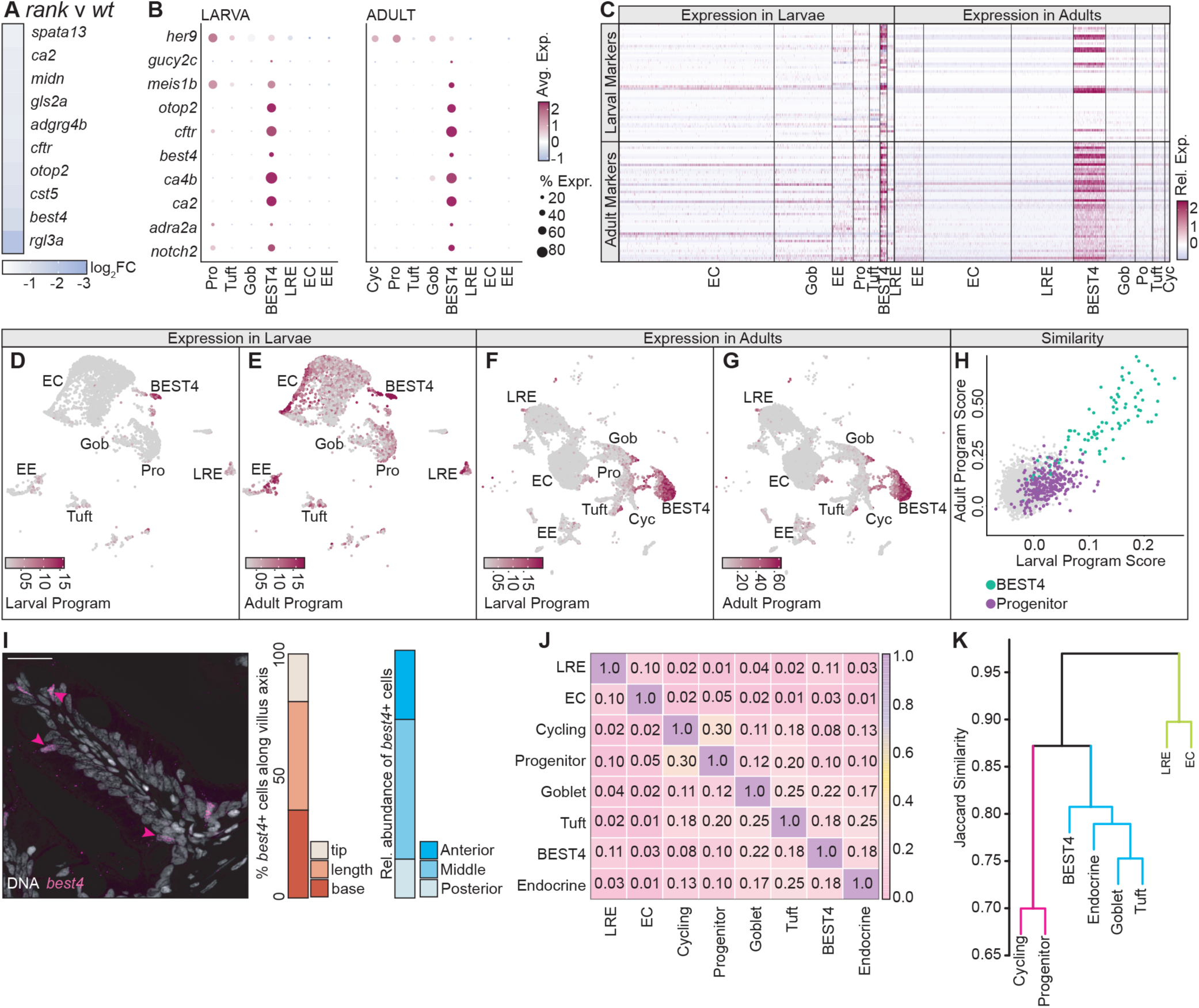
Comparative Analysis of BEST4 cells in the Adult Intestine. **A-B:** Expression of BEST4 cell marker genes in *rank* mutants relative to wildtype siblings (A), and cell-type expression patterns of established BEST4 cell markers in larval and adult intestines (B).**C:** Heatmap showing the relative expression of the top 50 larval and adult BEST4 cell markers in the intestinal epithelium of larvae and adults. **D-G:** UMAP projections of larval (D and F) and adult (E and G) epithelial cells colored by module scores for the larval BEST4 transcriptional program (D-E) and for the adult BEST4 transcriptional program (F-G). **H:** Scatter plot showing larval and adult BEST4 program scores for individual cells. BEST4 cells exhibit high scores for both programs, while progenitor populations show modest enrichment for BEST4 transcriptional programs. **I:** Representative fluorescence *in situ* hybridization image of *best4*+ cells (magenta) in the adult zebrafish intestine. Bar charts show the relative distribution levels of BEST4 cells along the villus axis (red shades), and the relative abundances of *best4+* cells in the anterior, middle, and posterior segments (blue shades). **J-K:** Jaccard similarity analysis of genes expressed in the respective cell types. **Abbreviations:** Pro: progenitor; Gob: Goblet; Cyc: cycling cells; LRE: lysosome-rich enterocytes; EC: enterocyte; EE: enteroendocrine.

To date, zebrafish BEST4 cells have been primarily studied in larvae, with minimal characterization in the adult intestine, an organ that more closely resembles the mammalian gut in terms of structural complexity and cellular composition. To better understand how BEST4 cells in the developing larval intestine relate to those in a mature adult intestine, we compared the gene expression profiles of larval and adult BEST4 cells in greater detail. Looking at the top 50 BEST4 cell marker genes from both developmental stages, we found that 24% of the markers were expressed at both stages and that those genes remained effective markers of the BEST4 cell populations (Figure 6C), suggesting partial conservation of genetic markers across developmental stages.

To better compare transcriptional programs between the developmental stages, we then identified the top 200 marker genes for BEST4 cells in larval and adult datasets. Each gene set was then treated as a transcriptional program and scored across cells using Seurat’s module scoring approach, allowing us to identify lineages that express larval and adult BEST4 programs in both datasets. As expected, the larval BEST4 gene program was largely restricted to larval BEST4 cells (Figure 6D), while the adult BEST4 program was predominantly restricted to adult BEST4 cells (Figure 6G), confirming that both gene sets effectively capture BEST4 cell identity at their respective stages. We next determined if these transcriptional programs are conserved across developmental stages. Here, we found that the larval BEST4 program strongly marked adult BEST4 cells (Figure 6F, H), indicating that key components of the larval transcriptional program persist in adult BEST4 cells. In contrast, the adult BEST4 program was not exclusively restricted to larval BEST4 cells (Figure 6E, H) but instead showed moderate activation across additional epithelial cell types. Together, these results suggest that adult BEST4 cells retain the core larval transcriptional program while acquiring additional gene expression programs that emerge during epithelial maturation.

To characterize adult BEST4 cells in greater detail, we then used fluorescence *in situ* hybridization to determine the rostro-caudal and apico-basal distribution of BEST4 cells throughout the adult intestine (Figure 6I). We discovered that BEST4 cells were quite evenly distributed along the villus axis, including at the villus base (Figure 6I). Our observation that *best4*+ cells exist at the base of fish villi contrasts with the predominately apical accumulation of BEST4 cells observed in human samples(1,36), and may reflect the fact that fish intestines lack crypt-like structures. We observed substantial differences for BEST4 cell distribution along the rostro-caudal axis. Although the frequency varied between samples, we consistently detected low frequency of BEST4 cells in the posterior intestinal segment, where *best4*+ cells accounted for approximately 2% of all intestinal epithelial cells, relative to 3.5% in the anterior segment and 7.1% in the middle intestinal section. (Figure 6I). We consider regional differences in BEST4 cell abundance interesting give the regionalized specialization of BEST4 cell functions observed in larval zebrafish and human tissues(35,37).

While BEST4 cells were initially classified as enterocyte-like absorptive cells(1), their relationship to other intestinal cell types remains unclear, with more recent work demonstrating that zebrafish BEST4 cells share developmental relationships with cells commonly considered “secretory”, including goblet, enteroendocrine and tuft cell lineages(35). Using our adult single cell gene expression data, we systematically compared the gene expression profiles of all adult epithelial cell types using Jaccard similarity, providing a quantitative measure of their relatedness (Figure 6J-K). Through pairwise comparisons, we identified three transcriptional branches: immature progenitor and cycling cells; metabolically active enterocytes and lysosome-rich enterocytes (LREs); and a distinct branch that encompassed BEST4 cells along with goblet, tuft, and enteroendocrine lineages, suggesting that BEST4 cells may belong to the “secretory” family of cells (Figure 6J). Our positioning of BEST4 cells within the secretory branch matches the observation that rat tuft and BEST4 cells share a common progenitor(38) and is supported by recent evidence that zebrafish BEST4 cells develop from *atoh1a*+ secretory cell progenitors(35).

### Interactions Between BEST4 cells and *Vibrio cholerae* in the Adult Intestine

BEST4 cells express prominent genes involved in immune regulation, including *ccl25a*, *cxcr4b* and *ptger4c* (Figure S6), and BEST4 cells are sensitive to diarrhea-causing bacterial toxins in human intestinal organoids(39). However, there are no *in vivo* data on the sensitivity of BEST4 cells to diarrheagenic pathogens. Therefore, we tested the effects of a *Vibrio cholerae* (*Vc*) infection on BEST4 cell populations in the adult intestine. We selected *Vc*, as zebrafish are susceptible to natural infections with *Vc*(40), and BEST4 cells are marked by expression of the ion channel *cftr* that initiates the massive chloride efflux responsible for diarrheal disease in cholera. To determine the impact of *Vc* on BEST4 cells, we enumerated *cftr*-positive cells in intestines of zebrafish that we challenged with *Vc* for 24h, followed by a one- or five-day recovery period (Figure 7A-C). In these experiments, we discovered that exposure to *Vc* led to a significant increase in the number of *cftr*-positive intestinal epithelial cells at both time points (Figure 7D), establishing an *in vivo* sensitivity of BEST4 cells to diarrheagenic bacteria. Notably, we also observed enhanced intestinal colonization by *Vc* in the intestines of *rank* mutant fish relative to their wildtype siblings at five days post-infection (Figure 7E). Combined, our data suggest that BEST4 cells are a widespread, pathogen-sensitive cell type in the adult zebrafish intestinal epithelium.

**Figure 7.**
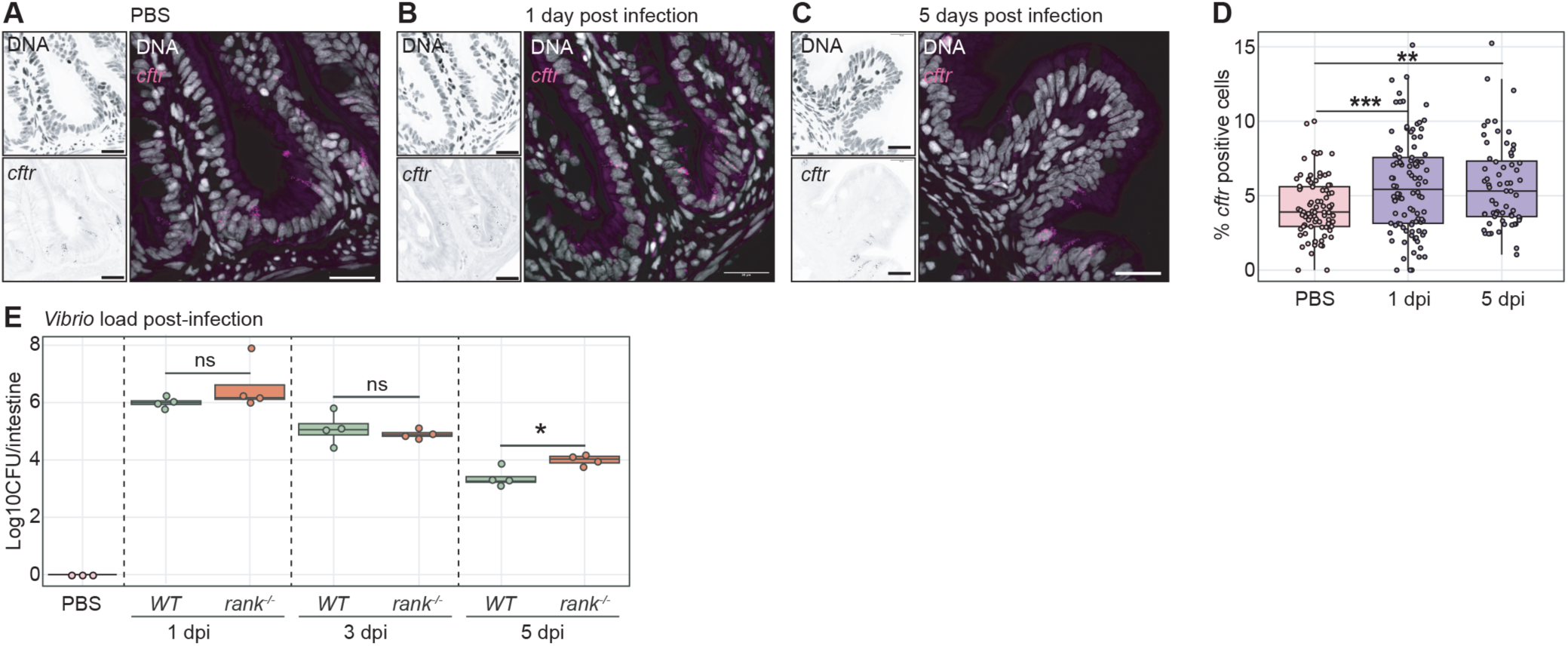
Interactions Between BEST4 Cells and *Vibrio cholerae* in the Adult Intestine. **A-C:** Fluorescence *in situ* hybridization images of sagittal intestinal sections from wildtype adult zebrafish showing the expression of *cftr* (magenta) in control fish (A), or *Vc*-infected fish that were give one (B) or five (C) days to recover from infection. **D:** Quantification of *cftr*-positive cells in intestines of the indicated treatment groups at the indicated numbers of days post-infection (dpi). Significance calculated by ANOVA with Tukey’s correction. **E:** Quantification of *Vibrio* burden in intestines of the indicated treatment groups at the indicated numbers of days post-infection (dpi). At each timepoint significance was calculated by an ANOVA. In each panel, one asterisk = p<0.05 two asterisks = p<0.01, and three = P < 0.001. In each image, scalebars are twenty micrometers, and arrowheads point to cells that express the gene of interest.

### RANK Contributes to BEST4 Cell Development

As RANK deficiency impacted expression of multiple BEST4 cell markers, we examined the effects of *rank* mutation on BEST4 cells. The 2F11 antibody labels intestinal epithelial cells positive for Annexin A4 (Anxa4), a gene that is expressed in the BEST4/secretory cell population (Figure 8A), including *pdia2*+ and *alox5a*+ tuft cells (Figure S7). We used immunofluorescence to confirm the existence of Anxa4-positive cells throughout the intestinal epithelium of *rank:GFP* adults, including in GFP-positive cells (Figure 8B), and we showed that loss of RANK significantly diminished the number of Anxa4-positive cells along the villus axis (Figure 8B-C), confirming a role for RANK in the development of Anxa4-positive cells. As Anxa4 marks multiple mature cell types, we systematically determined the consequences of RANK deficiency for goblet cells, enteroendocrine cells, and BEST4 cells throughout the intestinal epithelium.

**Figure 8.**
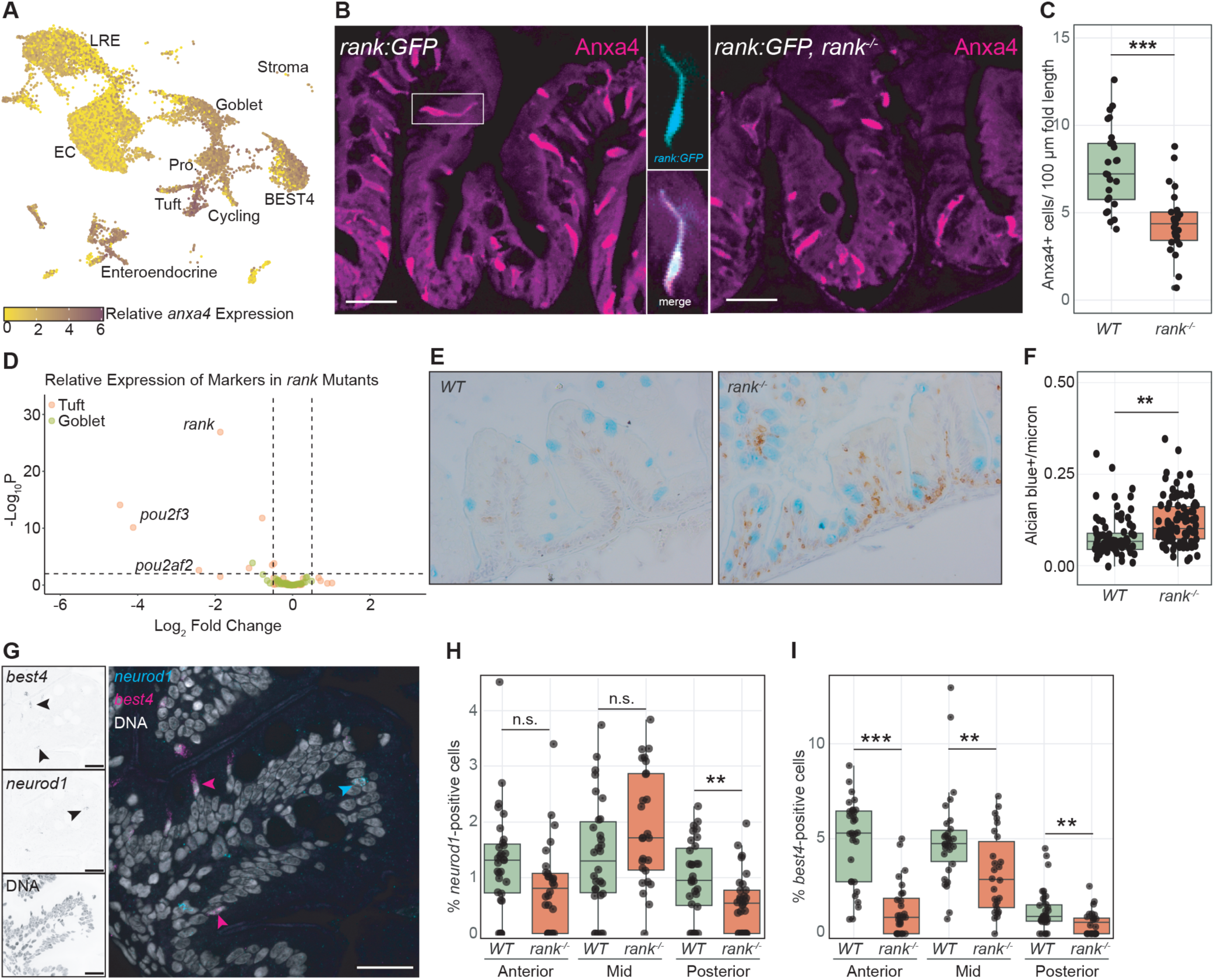
RANK Deficiency Leads to Epithelium-Wide Decrease in BEST4 Cell Numbers. **A:** Feature plot showing the relative expression of the 2F11 antibody target *anxa4* in the adult intestine. **B:** Visualization of Anxa4+ positive cells (magenta) in wildtype or RANK-deficient *rank:GFP* adult fish. Boxed region indicates a single GFP, Anxa4 double-positive cell in the merged image. **C:** Quantification of Anxa4-positive cells in wildtype and *rank* mutant intestines. **D:** Volcano plots showing relative differential expression of the top 20 goblet cell (sage) and tuft cell (apricot) markers in *rank* mutants relative to wildtype controls. **E-F:** Visualization (E) and quantification (F) of Alcian blue-positive goblet cells in wildtype and *rank* mutant adults. Both images were counterstained with the leukocyte marker Lcp-1 (brown) to assist orientation of samples. **G:** fluorescence *in situ* hybridization images of sagittal intestinal sections from wildtype fish showing pseudo colored expression of the BEST4 cell marker *best4* and the enteroendocrine cell marker *neurod1* in a representative wildtype intestine. **H-I:** Quantification of *neurod1* (H) and *best4* (I)-positive epithelial cells in the anterior, middle, and posterior intestinal segments of wildtype and *rank* mutant intestines as indicated. In all panel, two asterisks = P< 0.01, and three = P < 0.001, calculated by ANOVA. In each image, scalebars are twenty micrometers, and arrowheads point to cells that express the gene of interest.

Loss of RANK did not affect expression of any of twenty goblet cell marker genes (Figure 8D) or lower the number of intestinal epithelial goblet cells (Figure 8E). Instead, we observed a significant increase in the number of Alcian blue-positive goblet cells in *rank* mutants, relative to their wildtype sibling controls (Figure 8E-F), indicating that RANK deficiency enhances goblet cell specification without an increase in the global expression of goblet cell markers, suggesting a possible decline in marker expression per goblet cell in *rank* mutants. We consider the inverse relationship between goblet and BEST4 cell numbers in *rank* mutants interesting, given reports that BEST4 and goblet cells are often found in pairs(41,42), and that treatment of mouse small intestinal organoids with RANK ligand suppresses expression of goblet signature genes(43). Unlike tuft and goblet cells, RANK deficiency only affected the representation of enteroendocrine cells in the posterior intestinal epithelium, where it led to a significant loss of *neurod1+* enteroendocrine cells (Figure 8G-H). In contrast, RANK deficiency significantly impacted BEST4 cell numbers throughout the intestinal epithelium, leading to a greater than 50% drop in the BEST4 cell population in each section (Figure 8G, I).

Combined, our results uncover an epithelium-wide role for the zebrafish *rank*/*tnfrsf11a* ortholog in the development of intestinal epithelial type-two tuft cells and BEST4 cells, and a restricted requirement for RANK in the development of posterior enteroendocrine cells and anterior intestinal type one tuft cells.

## DISCUSSION

At the intestinal barrier, specialized cells harvest nutrients, protect from pathogens, and coordinate tolerant immune responses to the resident microbiota. As epithelial breaches fuel inflammatory disease, it is essential that we decipher the pathways responsible for barrier formation and maintenance. RANK has critical roles in the mammalian intestine, where it controls M cell development(11,13), epithelial remodelling during pregnancy(14), ILC3 activity(15), and antiparasitic defenses(16). However, we know less about effects of RANK deficiency on homeostatic maintenance of the gut epithelium. In this study, we determined the consequences of systemic RANK deficiency for the adult zebrafish gut. We found that loss of RANK diminished the epithelial abundance of immune-modulatory tuft cells and BEST4 cells and showed that RANK deficiency led to an accumulation of goblet cells and activated leukocytes, suggesting that RANK signaling modifies immune homeostasis in the zebrafish gut. As a caveat, we note that our study detailed the consequences of systemic RANK deficiency for the fish gut. Thus, our work does not address cell-autonomous roles for RANK signaling in the specification of individual cell types, or in the regulation of intestine immune homeostasis. Moving forward, we feel our results raise several questions that merit investigation.

How do zebrafish tuft cells relate to mammalian equivalents? As zebrafish intestines contain ILC2s and ILC3s(44), and fish encode immune-regulatory *il-4*, *il-13*, and *il-10* orthologs(45,46), we view zebrafish as an excellent animal to characterize type two mucosal immunity. Mammalian tuft cells are a relatively rare post-mitotic lineage that activates antiparasitic defenses(47), contributes to antibacterial immunity(7), and provides a reserve population of stem cells for injury-dependent tissue regeneration(9). Recent work indicated that mammalian tuft cells arise from a crypt-associated progenitor cell that transitions through tuft-1 and tuft-2 maturation states as it migrates apically. We discovered that, like mammals, zebrafish intestines contain two *pou2f3*-positive tuft cell subtypes, including one marked by expression of receptors for IL-4 and IL-13, and genes required for production of pro-inflammatory leukotrienes, indicating the existence of specialist fish tuft cells dedicated to type two defenses. We do not know if the fish Tuft-1 and Tuft-2 cell types represent functionally distinct tuft cell subsets, or if they represent maturation states like those observed in mice. Our imaging data indicate that both cell types are found along the villus axis, and we rarely observed co-expression of *pdia2* and *alox5a* in individual cells. Furthermore, our gene expression trajectory analysis suggests that both cell types correspond to distinct terminal maturation states. Combined, we feel these data support a model that both subsets are functionally distinct. However, precise lineage labeling experiments are essential to fully test this hypothesis.

Our data also point to differences between immune effector functions in fish and mammalian tuft cells. In mice, tuft cell-derived IL-25 is essential for tuft cell-dependent induction of type two immunity(5–7,48,49). As the fish genome does not appear to encode an *il-25* ortholog, it is unclear if, or how, fish tuft cells engage neighboring ILC2s. However, intestinal tuft cells from humans and non-human primates also do not seem to express IL-25(50,51). Thus, we consider it possible that additional mediators of tuft cell-ILC2 communication require identification. Perhaps the most striking difference between fish and mammalian tuft cells is the apparent absence of *choline acetyl transferase* expression in fish tuft cells. In mammals, acetylcholine is a prominent effector of tuft cell-dependent immunity(52), including direct effects on helminth fecundity(53), suggesting that fish tuft cells may rely on a distinct class of effectors to protect from infection. In the future, it will be of interest to determine the impacts of fish tuft cells on antiparasitic immunity in live infection models.

What is the relationship between tuft cells, BEST4 cells, and goblet cells? We often view development of mature intestinal epithelial cells as a metronomic progression through maturation states that are controlled by germline-encoded signaling pathways. However, epithelial maturation is a more stochastic event that is also swayed by damaging agents, pathogen-derived virulence factors, or metabolic products of commensal bacteria. For example, luminal parasites drive a rapid accumulation of tuft and goblet cells that initiate the “weep and sweep” elimination of the parasite(4–6), while commensal bacteria promote expansion of goblet cell numbers and a thickening of the mucus barrier(54,55). Our work suggests that BEST4 cells are part of this plastic community of microbe-sensitive intestinal epithelial cells. From a transcriptional perspective, we find that BEST4 cells are most like secretory cells with the closest relationship to goblet and tuft cells, and we present evidence that RANK deficiency affects the epithelial abundance of type two tuft cells, BEST4 cells, and goblet cells. Our findings match data that tuft and BEST4 cells share a common progenitor in rats(38), and align with reports that exposure of mouse small intestinal organoids to RANK ligand enhances expression of tuft signature genes(16), while suppressing expression of goblet cell signatures(43). In agreement with data from human intestinal organoids(39), we present *in vivo* evidence that *Vibrio cholerae* promote accumulation of BEST4 cells in the fish gut, and we demonstrate that RANK deficiency impairs clearance of *Vibrio*, supporting a role for BEST4 cells in sensing and responding to luminal pathogens. In the future, it will be valuable to identify the factors responsible for development of BEST4 and secretory lineages in the presence or absence of pathogenic challenges.

How do BEST4 cells relate to M cells? In mammals, RANK is essential for M cell development(11,56). As fish intestines do not appear to include M cells, we cannot chart the developmental relationships between M cells and BEST4 cells in the fish gut. However, our finding that RANK deficiency leads to a loss of BEST4 cells matches a report that human BEST4 cell development proceeds through a transient M-like cell state(1). In contrast, recent work with human intestinal organoids showed that treatment with RANK ligand led to a loss of BEST4 cells, and a concomitant gain of M cells, indicating differential effects of RANK signaling on BEST4 and M cell development(39). However, we feel it important to note that organoid growth media do not typically permit simultaneous development of BEST4 and M cells. As a result, it is challenging to extrapolate these data to an *in vivo* appreciation of the factors that ultimately balance BEST4 and M cell specification in the intestine.

Are BEST4 cells involved in intestinal epithelial defenses? First described in a histological study of human intestinal epithelial cells(41), BEST4 cells were subsequently profiled in single cell gene expression atlases that showed BEST4 cells express ion channels, such as *best4* and *cftr*, alongside carbonic anhydrases needed for acid-base homeostasis, pointing to possible roles for BEST4 cells in mucus hydration and luminal pH(1,2,36,50,57,58). At the same time, patient data implicated BEST4 and OTOP2(59,60) mutations in poor colorectal cancer prognosis(61), underscoring a need to understand how BEST4 cells function *in vivo*. Since their identification in humans, BEST4 cells have been found in zebrafish(23,24,33,34), macaques(62), pigs(63), rabbits(32), pythons(64), and rats(38), and a recent study identified a core transcriptional program in most BEST4 cells(35), suggesting an evolutionarily conserved role for BEST4 cells in gut function. Given their physical proximity to goblet cells, their putative roles in pH regulation and mucus hydration, and their sensitivity to enteric pathogens, we hypothesize that BEST4 cells may contribute to the regulation of host-microbe interactions in the gut. In this regard, we consider it particularly intriguing that zebrafish BEST4 cells express elevated levels of the prostaglandin *ptger4c* receptor(34), as well as the *ccl25a* chemokine (24). As mice intestines lack BEST4 cells, we believe zebrafish are an ideal animal model to explore roles for BEST4 cells in host responses to commensal and pathogenic microbes.

### Limitations of our study

As we examined the consequences of systemic RANK deficiency for cell composition and function in the intestinal epithelium, our work does not allow us to unambiguously determine the source of RANK signals, or to identify the key responding cells. Prior work showed that tuft and goblet cells are sensitive to RANK exposure in organoids(16,43), suggesting that the epithelium is capable of direct responses to RANK signaling. However, several studies have identified important roles for immune cell-derived RANK in the coordination of epithelial cell behavior(65–68). Our single cell atlas suggests minimal expression of *rank* in gut-associated leukocytes(24). Nonetheless, we feel it will be important to examine the consequences of leukocyte-specific RANK ablation for epithelial cell dynamics in future studies. Furthermore, we note that our study characterized intestinal physiology in a *rank* mutant that simultaneously affected the epithelial representation of three cell types: tuft cells, BEST4 cells and goblet cells. As a result, our work does not allow us to make any inferences related to the developmental relationships of the respective cell types. For example, it is possible that BEST4 and tuft cells share a common RANK-sensitive progenitor, but it is equally possible that loss of one cell type indirectly affects the abundance of the other. Moving forward, it will be of interest to identify the signaling networks responsible for the developmental specification of the respective lineages.

## MATERIALS AND METHODS

### Animal Husbandry and zebrafish lines

All zebrafish experiments were approved by the University of Alberta’s Animal Care & Use Committees of Biosciences and Health Sciences (AUP#3032), operating under the guidelines of the Canadian Council of Animal Care. Wild type TL, Tg(*rank:GFP*), and Tg(*NFkB:GFP*) were reared at 27°C in facility water (Instant Ocean Sea Salt dissolved in reverse osmosis water maintained at a conductivity of 1000µS and pH buffered to pH 7.0 with sodium bicarbonate) under a 14h/10h light/dark cycle using standard zebrafish husbandry protocols. Adult zebrafish were fed GEMMA Micro 300 daily at 8AM, followed by live rotifers daily at 2:30PM. Embryos raised to larvae were housed at 29°C under a 14h/10h light/dark cycle.

### Generating rank mutants

Tg(*rank*:GFP) was described previously(23). To generate rank mutants, CRISPR-Cas9 mutagenesis was carried out as described previously(69) using the IDT AltR CRISPR-Cas9 System. Briefly, the cytoplasm of single-celled Tg(*rank:GFP*) embryos were injected with approximately 1nL of 5µM gRNA:Cas9 ribonucleoprotein complex: duplexed gRNA, trans-activating crispr RNA (tracrRNA; IDT 1072533), and Cas9 enzyme (IDT) in IDT duplex buffer (IDT 11-05-01-03). To generate stable mutants, injected embryos were reared to adulthood, then single adults were outcrossed to AB wild-type fish. F1 progeny were reared, in-crossed and genotyped by fin clipping to search for F2 generations bearing a frame-shift mutation. PCR amplicons were generated with the following primers: *rank* diagnostic forward primer CACACACGCACATGAACTATAACC; *rank* diagnostic reverse primer GTCTTAAAGTGACACGAACCC. Cutting efficiency was assessed by PCR and amplicon digestion with the restriction enzyme MslI (NEB R0571L), where lack of digestion indicated disruption of the recognition sequence. Amplicons were sequenced at the Molecular Biology Services Unit at the University of Alberta to determine the nature of the lesion.

### Generating rank crispants in Tg(*NF-kB*:EGFP) background

To investigate the effect of *rank* activity on NF-kB:EGFP, single cell-stage NF-kB:EGFP embryos were injected with Cas9 enzyme (control, no sgRNA) or Cas9 enzyme complexed with *rank* sgRNA to generate *rank* crispants. At 6 days post fertilization, the larvae were euthanized with tricaine, fin clipped, genotyped and fixed overnight at 4°C. Larvae were washed three times with PBS then embedded in 0.7% UltraPure low melting point agarose (Invitrogen Cat#16520) on a glass bottom dish.

### HCR™ RNA Fluorescence *in situ* hybridization

Molecular Instruments Multiplexed HCR™ RNA-FISH (v3.0) protocol for Fresh/Fixed Frozen Tissues was followed to detect mRNA of interest in adult intestinal tissue sections (5µm thickness) and has been described previously(24). HCR-RNA FISH probes used in this paper were *alox5a*, *best4*, *cftr*, *her15.1*, *neurod1*, *pdia2*, *pou2f3*, and *tnfrsf11a* (*rank*).

### Immunofluorescence

We used previously established morphological criteria(31) to identify the anterior (rostral intestinal bulb), middle (mid-intestine), and posterior (caudal intestine) intestinal segments. Zebrafish intestines were fixed in 4% paraformaldehyde (diluted in PBS from a 16% stock: EMS Cat#15710) overnight at 4°C. Intestines were washed twice in 1xPBS then cryoprotected in 15% (w/v) sucrose/PBS at room temperature until sunk followed by sinking in 30% (w/v) sucrose/PBS (overnight at 4°C). Intestines were mounted in optimal cutting temperature embedding medium (Fisher Scientific Cat #23-730-571), then frozen on dry ice. Five micrometer sections were collected on Superfrost Plus slides (Fisherbrand Cat #22-037-246). After allowing sections to adhere onto slides, slides were immersed in PBS for 20 minutes at room temperature. Tissue was blocked for 1h at room temperature in 3% (w/v) Bovine Serum Albumin dissolved in PBST (1xPBS + 0.2% (v/v) Tween-20). Primary antibodies were diluted in blocking buffer then layered onto sections. Sections were incubated in primary antibody solution overnight in a humid chamber at 4°C. Excess primary antibody was washed by immersing slides in fresh PBST for 1.5h. Sections were incubated in secondary antibody solution for 1h at room temperature in a humid chamber, protected from light. All secondaries were prepared at a 1:2000 dilution in blocking buffer; Phalloidin was added to the secondary antibody solution at 1:500. Secondary solution was removed then sections incubated in Hoechst diluted 1:2000 in PBST for 10 minutes protected from light. Slides were washed in PBST by brief immersion followed by a 30-minute incubation in fresh PBST protected from light. Slides were mounted in Fluoromount™ Aqueous Mounting Medium (Sigma F4680-25mL). Primary antibodies used for immunofluorescence in this study: Chicken polyclonal anti-GFP (ThermoFisher Scientific Cat# PA1-9533) at 1:1000; Mouse anti-zebrafish gut secretory cell epitopes (abcam Cat# ab71286) at 1:500. Fluorescent secondaries and stains used in this study were: Goat anti-chicken IgY (H+L) secondary antibody, Alexa Fluor™ 488 (ThermoFisher Scientific Cat# A-11039); Goat anti-Rabbit IgG (H+L) Cross-Adsorbed Secondary Antibody, Alexa Fluor™ 568 (ThermoFisher Scientific Cat# A-11011); Alexa Fluor™ 647 Phalloidin (ThermoFisher Scientific Cat# A22287); Hoechst 33258 (Molecular Probes Cat #H-3569).

### Lcp1 Immunohistochemistry & Alcian blue staining

Heterozygous Tg(*rank:GFP*); *rank* +/- were in-crossed and progeny co-housed. At 7 weeks post fertilization, the progeny were genotyped then five fish per genotype (rank +/+ and rank -/-) were co-housed until 10 weeks post fertilization. Experimental fish were fasted for 24h prior to dissection. Whole intestines from 10-week-old fish were dissected then fixed overnight at 4°C in BT fixative (4% PFA, 0.15mM CaCl2, 4% sucrose in 0.1M phosphate buffer pH 7.4). Whole intestines were processed for paraffin embedding at the University of Alberta Biological Sciences Advanced Microscopy Facility then IHC performed on 5µm tissue sections as described previously(24). L-Plastin was detected in 10-week-old intestines using rabbit anti-Lcp1 antibody (GeneTex GTX124420, 1/10000) followed by colorimetric detection using Cell Signaling Technology® SignalStain® Boost Detection Reagent HRP Rabbit (Cat 8114) and SignalStain® DAB Substrate Kit (Cat 8059, 10 minutes). Slides were then stained with Alcian blue for 3 minutes followed by a 10-minute rinse in running tap water. Nuclei were counterstained with ¼-strength hematoxylin (30 seconds) followed by another 10-minute rinse in running tap water. Tissue was then dehydrated, cleared with toluene, and mounted in Dpx. The fish used in this experiment were also used to measure body length and whole intestine length.

### Microscopy

Histological samples were imaged on a ZEISS AXIO A1 compound light microscope with a SeBaCam 5.1MP camera. Confocal images of intestinal tissue sections were captured on an Olympus IX-81 microscope fitted with a CSU-X1 spinning disk confocal scan-head, Hamamatsu EMCCD (C9100-13) or Hamamatsu Orca-Fusion BT digital sCMOS camera and operated with PerkinElmer’s Volocity. Whole larval confocal imaging was performed on a Leica Falcon SP8 equipped with a 25x 0.95NA Water HC Fluotar objective lens then stitched with Leica Application Suite X software. Fiji(70) was used to generate maximum intensity projections of confocal Z-stacks and false color manipulations.

### Bulk RNAseq of whole intestines

Heterozygous Tg(*rank:GFP*); rank +/- were in-crossed and progeny co-housed. At 7 weeks post fertilization, the progeny were genotyped then six fish per genotype (rank +/+ and rank -/-) were co-housed until 8 weeks post fertilization. Experimental fish were fasted for 24h prior to dissection for RNA extraction. Whole intestines from 8-week-old fish were dissected then homogenized in Trizol Reagent then RNA was extracted following manufacturer’s suggestions. Fish carcasses were fin clipped and genotyped by Sanger sequencing. Pure RNA was sent to Novogene for polyA enrichment of the RNA library (unstranded) and sequencing on NovaSeq X Plus (PE150, Q30≥85%)(6Gb raw data per sample).

### Bulk RNA-Seq quality control and alignment

The fastq files were processed using the default parameters of the nf-core RNA-seq pipeline (version 3.9)(71,72). Quality control metrics were assessed via FASTQC (version 0.11.9). Adaptors and low-quality reads were filtered out using the default parameters of TrimGalore! (version 0.6.7). STAR (version 2.7.10a) was used to map reads to the zebrafish genome (GRCz11). Salmon (version 1.5.2) quantified transcripts(73).

### Bulk RNA-Seq data processing and DESeq2 analysis

The quantification files produced by Salmon were used for further analysis with the DESeq2 (version 1.38.3) R package(74), which also assigned Zebrafish Information Network symbols to each gene. Differential expression analyses were completed with DESeq2 to identify genes differentially expressed between wild type and *rank^-/-^*fish, with an adjusted *p*-value < 0.01 considered significant. Significantly differentially expressed genes were then used for principal component analysis, which was also completed with DESeq2.

### Single cell gene expression analyses

All single cell gene expression work was done in R with previously published larval and adult data sets(23,24). Scripts for identifying individual clusters, identifying marker gene expression, and for pseudotime mapping of tuft cell development are available at https://github.com/rjwllms/Thesis-scripts. All GO term analysis was performed using DAVID gene ontology analysis(75,76). To compare BEST4 transcriptional programs between developmental stages, the top 200 markers from larval and adult BEST4 cells were used to define stage-specific transcriptional programs. These gene sets were scored across cells in both datasets using Seurat’s AddModuleScore function, which calculates an aggregated expression score for each gene set while controlling for gene expression background. Program scores were then visualized on UMAP embeddings and compared across cell populations to assess the distribution of larval and adult BEST4 transcriptional programs across developmental stages.

### Cross-species comparison of zebrafish and mouse tuft cells

Single-cell RNA sequencing datasets containing intestinal tuft cells from zebrafish and mouse were analyzed using the Seurat R package(77). To enable cross-species comparison, zebrafish genes were mapped to mouse orthologs using ortholog annotations obtained from Ensembl. When multiple orthologs were present, the first listed ortholog was retained to generate a one-to-one gene mapping table. Only genes with identifiable orthologs in both species were retained for downstream analysis. To quantify transcriptional similarity between zebrafish and mouse tuft cell subtypes, average gene expression profiles were computed for each cluster. Correlation coefficients were calculated between zebrafish and mouse cluster-level expression profiles using genes shared between species following ortholog mapping. These values were used to generate the cross-species correlation heatmap. For joint visualization of zebrafish and mouse tuft cells, ortholog-mapped gene expression matrices were merged and integrated using the Seurat integration workflow. To quantify the extent of cross-species mixing within each cluster, a nearest-neighbor graph was constructed from the integrated dataset using Seurat’s FindNeighbors function. For each cell, the fraction of nearest neighbors derived from the opposite species was calculated.

## ACKNOWLEDGMENTS

We are very grateful to Dr. Jeffrey Farrell at the NIH for constructive feedback on this manuscript, and to Dr. Jennifer Hocking at the University of Alberta for expert guidance on zebrafish handling. We would also like to thank the University of Alberta North Campus Animal Services for their excellent care of the zebrafish aquatics facilities at the University of Alberta. Imaging experiments were performed at the University of Alberta Faculty of Medicine & Dentistry (FoMD) Cell Imaging Core, RRID:SCR_019200, which receives financial support from FoMD, the Department of Medical Microbiology and Immunology, the University Hospital Foundation, and Canada Foundation for Innovation (CFI) awards to contributing investigators. Immunohistochemistry experiments were performed at the University of Alberta’s Faculty of Science Department of Biological Sciences Advanced Microscopy Facility. The Advanced Microscopy Facility is supported by Alberta Innovation and Science, Canadian Foundation for Innovation, National Science and Engineering Research Council, University of Alberta, and Faculty of Science. This work used computing resources provided by the Stanford Genetics Bioinformatics Service Center. This work was supported by a grant from the Canadian Institutes of Health Research (grant no. MOP77746) to EF. RJW received funding support through the National Science and Engineering Research Council Graduate Scholarships, and Alberta Innovates Graduate Student Scholarships.

## SUPPLEMENTARY FIGURES

**Figure S1.**
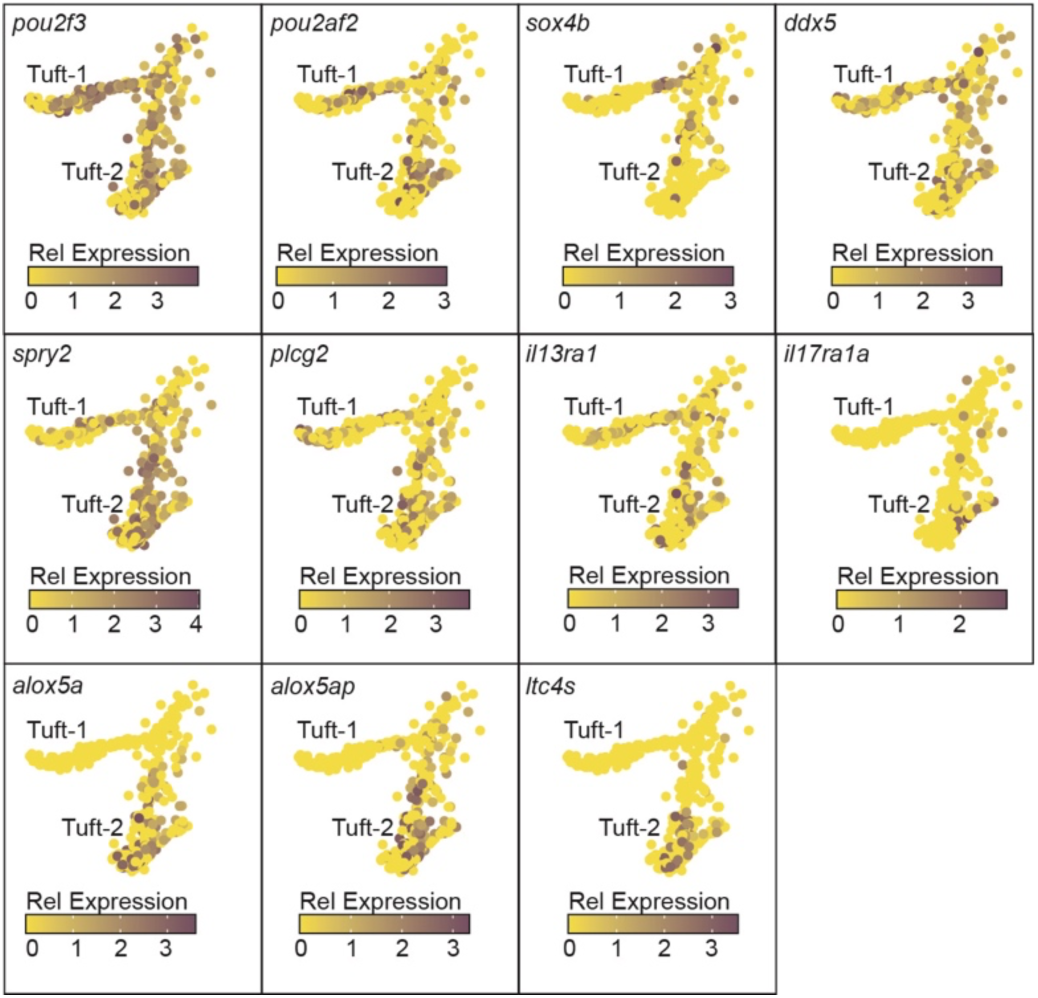
Expression of classical tuft cell markers (panels from *pou2f3* to *plcg2*) and immune regulatory markers (panels from *il13ra1* to *ltc4s*) in zebrafish tuft cells.

**Figure S2.**
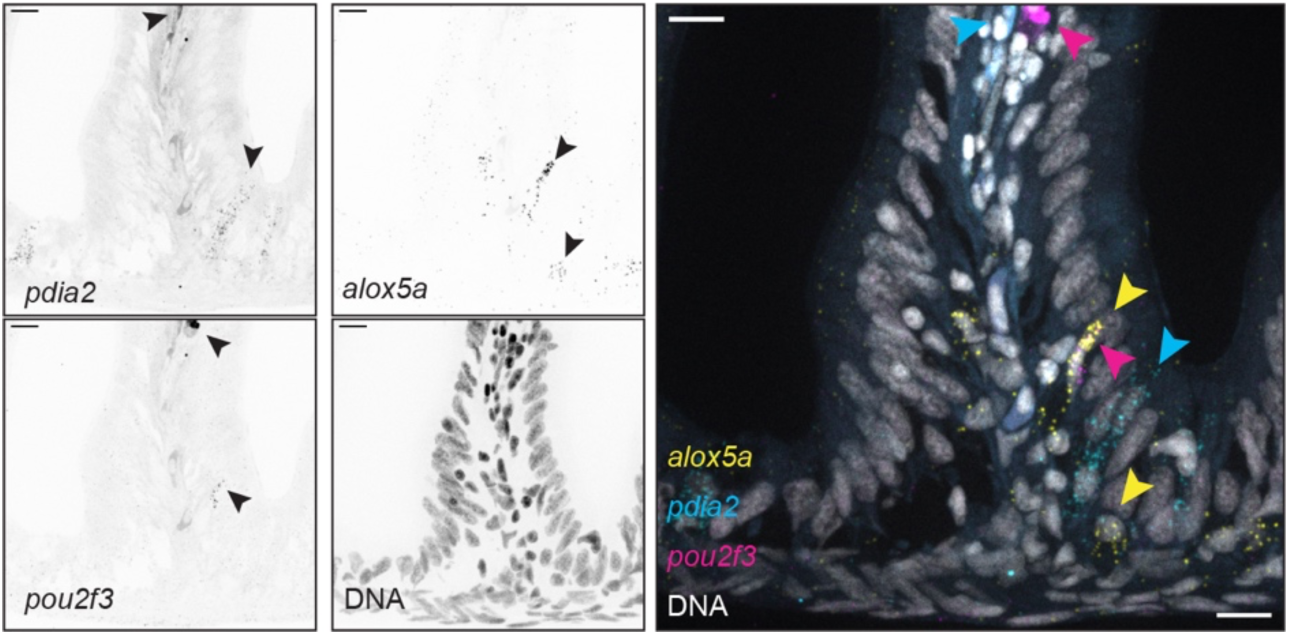
Fluorescence *in situ* hybridization showing the expression of the indicated genes in grayscale for each gene, and pseudo colored in the merged image. In each image, scalebars are twenty micrometers, and arrowheads point to cells that express the gene of interest.

**Figure S3.**
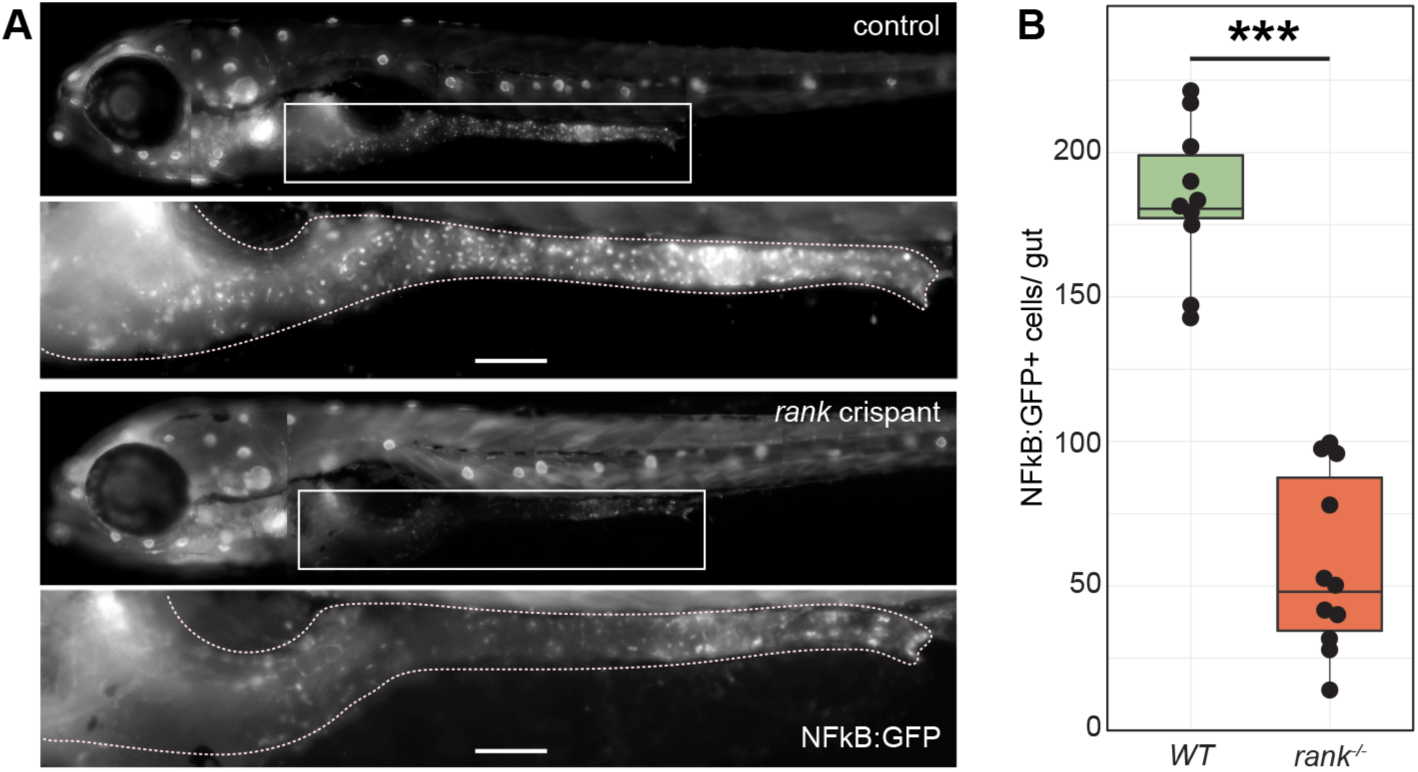
Visualization (A) and quantification (B) of GFP expression in control or *rank* crispant NF-kB:GFP 6 dpf larvae.

**Figure S4.**
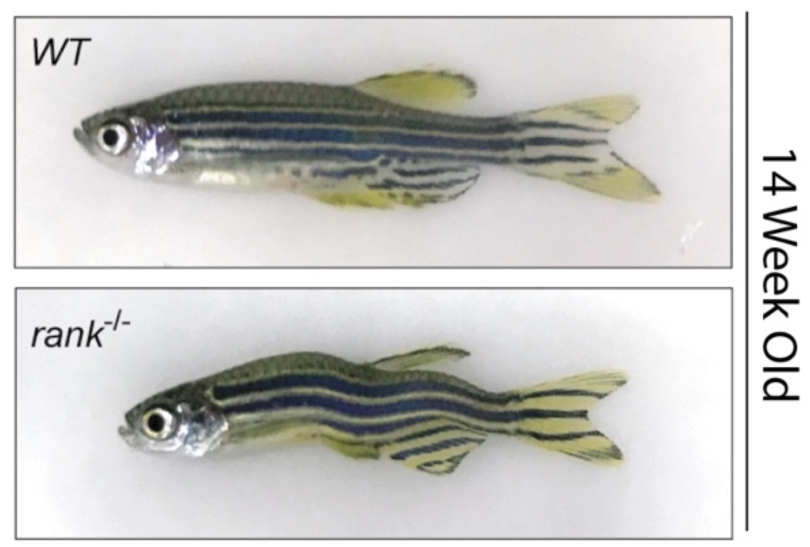
Spinal curvature defects in fourteen-week-old *rank* mutant fish relative to wildtype (WT) controls.

**Figure S5.**
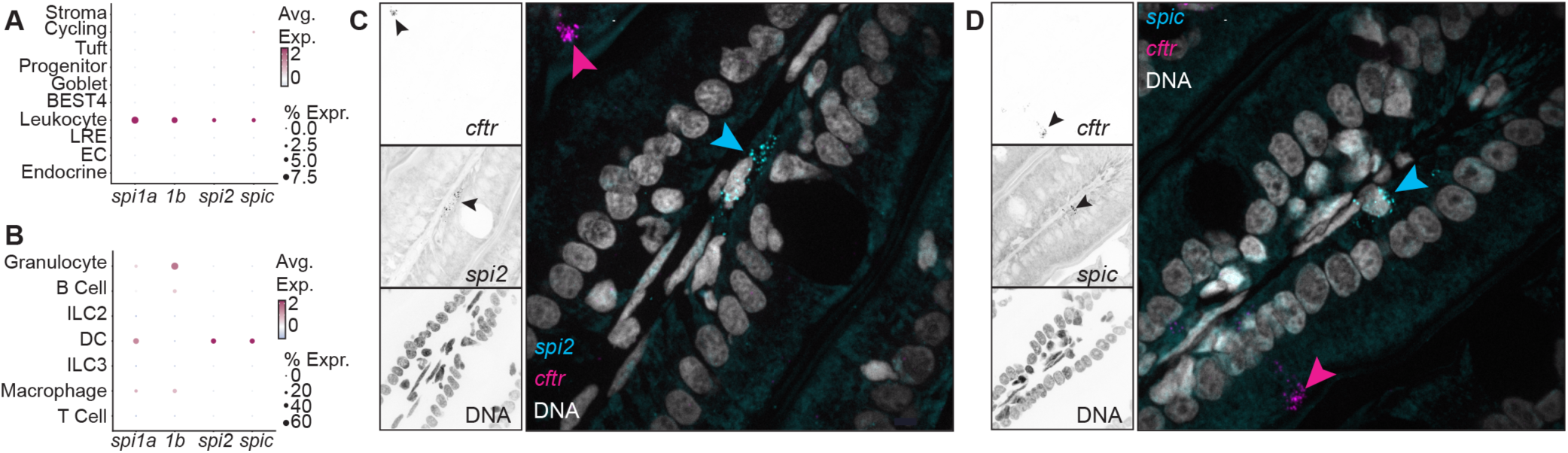
**A-B:** Expression patterns of zebrafish *spi* family members in the zebrafish guts (A) and leukocyte cell types (B). **C-D:** Visualization of *cftr* co-expression with *spi2* (C) and *spic* (D) in adult zebrafish intestines. Whereas *cftr* marks BEST4 cells, *spi2* and *spic* mark vasculature-associated dendritic cells.

**Figure S6.**
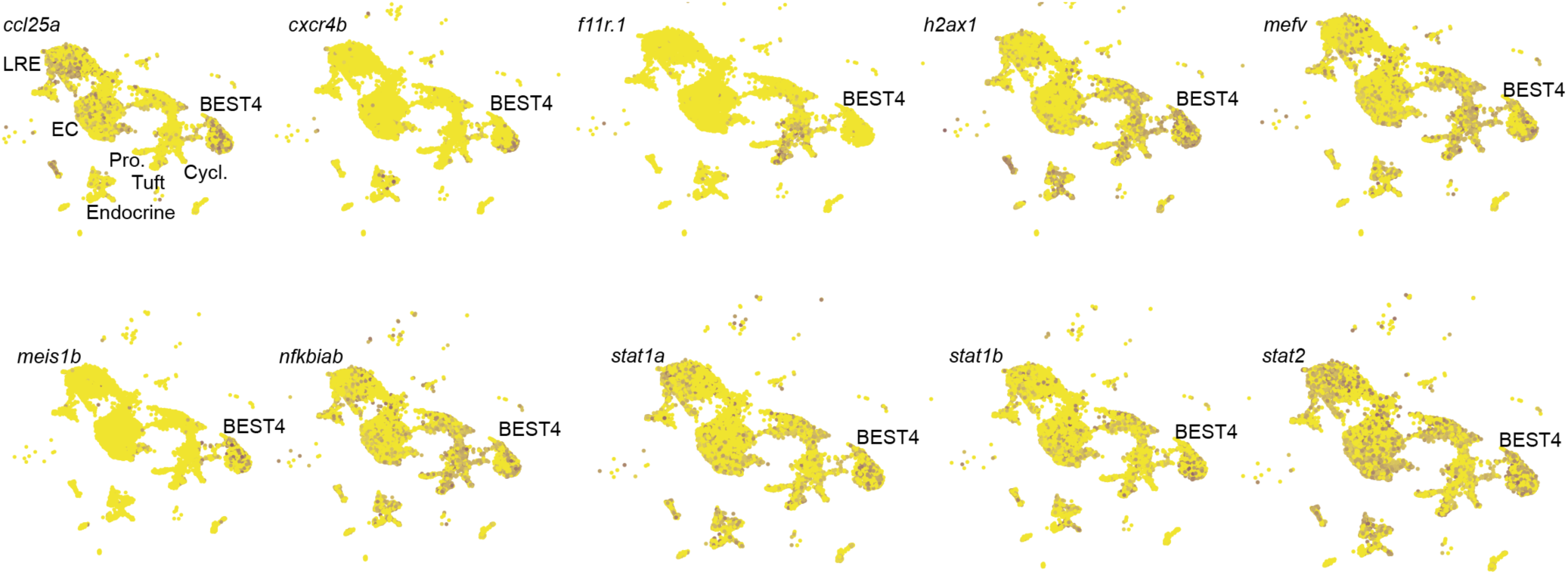
A series of feature plots showing expression of multiple immune regulatory genes in adult intestinal BEST4 cells.

**Figure S7.**
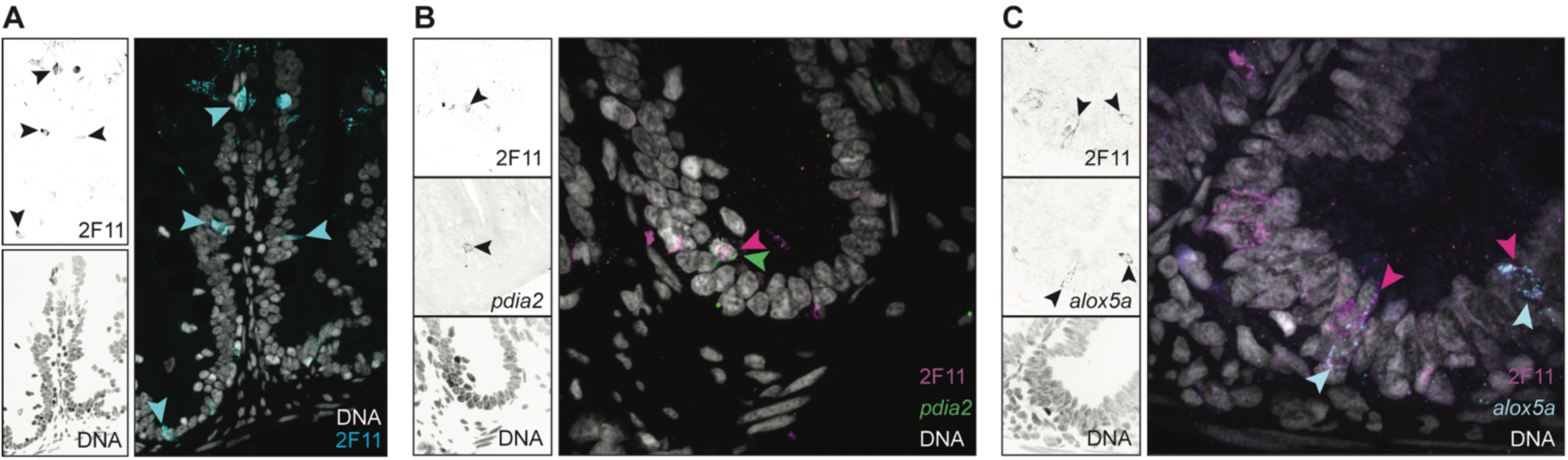
**A:** Visualization of 2F11+ cells (marked with arrowheads) in the intestinal epithelium of an adult zebrafish. **B-C:** Dual staining of intestines for 2F11 (B-C) and *pdia2* (B) or *alox5a* (C). In the false-colored images, DNA is labeled in grey, 2F11 is labeled in magenta C, *pdia2* is labeled green B, and *alox5a* is labeled cyan C. In each case, the tuft cell markers overlap with the 2F11 antibody that labels epithelial secretory cells.

## CITATIONS

1. Smillie CS, Biton M, Ordovas-Montanes J, Sullivan KM, Burgin G, Graham DB, et al. Intra- and Inter-cellular Rewiring of the Human Colon during Ulcerative Colitis. Cell. 2019 Jul 25;178(3):714–730.e22. doi:10.1016/j.cell.2019.06.029

2. Parikh K, Antanaviciute A, Fawkner-Corbett D, Jagielowicz M, Aulicino A, Lagerholm C, et al. Colonic epithelial cell diversity in health and inflammatory bowel disease. Nature. 2019 Mar;567(7746):49–55. doi:10.1038/s41586-019-0992-y

3. Feng X, Flüchter P, De Tenorio JC, Schneider C. Tuft cells in the intestine, immunity and beyond. Nat Rev Gastroenterol Hepatol. 2024 Dec;21(12):852–68. doi:10.1038/s41575-024-00978-1

4. Howitt MR, Lavoie S, Michaud M, Blum AM, Tran SV, Weinstock JV, et al. Tuft cells, taste-chemosensory cells, orchestrate parasite type 2 immunity in the gut. Science. 2016 Mar 18;351(6279):1329–33. doi:10.1126/science.aaf1648

5. Gerbe F, Sidot E, Smyth DJ, Ohmoto M, Matsumoto I, Dardalhon V, et al. Intestinal epithelial tuft cells initiate type 2 mucosal immunity to helminth parasites. Nature. 2016 Jan 14;529(7585):226–30. doi:10.1038/nature16527

6. von Moltke J, Ji M, Liang HE, Locksley RM. Tuft-cell-derived IL-25 regulates an intestinal ILC2–epithelial response circuit. Nature. 2016 Jan;529(7585):221–5. doi:10.1038/nature16161

7. Xiong Z, Zhu X, Geng J, Xu Y, Wu R, Li C, et al. Intestinal Tuft-2 cells exert antimicrobial immunity via sensing bacterial metabolite N-undecanoylglycine. Immunity. 2022 Apr 12;55(4):686–700.e7. doi:10.1016/j.immuni.2022.03.001

8. Eshleman EM, Rice T, Potter C, Waddell A, Hashimoto-Hill S, Woo V, et al. Microbiota-derived butyrate restricts tuft cell differentiation via histone deacetylase 3 to modulate intestinal type 2 immunity. Immunity. 2024 Feb 13;57(2):319–332.e6. doi:10.1016/j.immuni.2024.01.002

9. Huang L, Bernink JH, Giladi A, Krueger D, van Son GJF, Geurts MH, et al. Tuft cells act as regenerative stem cells in the human intestine. Nature. 2024 Oct;634(8035):929–35. doi:10.1038/s41586-024-07952-6

10. Ono T, Hayashi M, Sasaki F, Nakashima T. RANKL biology: bone metabolism, the immune system, and beyond. Inflamm Regen. 2020 Feb 7;40(1):2. doi:10.1186/s41232-019-0111-3

11. Knoop KA, Kumar N, Butler BR, Sakthivel SK, Taylor RT, Nochi T, et al. RANKL is necessary and sufficient to initiate development of antigen-sampling M cells in the intestinal epithelium. J Immunol Baltim Md 1950. 2009 Nov 1;183(9):5738–47. doi:10.4049/jimmunol.0901563

12. Nagashima K, Sawa S, Nitta T, Tsutsumi M, Okamura T, Penninger JM, et al. Identification of subepithelial mesenchymal cells that induce IgA and diversify gut microbiota. Nat Immunol. 2017 Jun;18(6):675–82. doi:10.1038/ni.3732

13. Kanaya T, Sakakibara S, Jinnohara T, Hachisuka M, Tachibana N, Hidano S, et al. Development of intestinal M cells and follicle-associated epithelium is regulated by TRAF6-mediated NF-κB signaling. J Exp Med. 2018 Jan 16;215(2):501–19. doi:10.1084/jem.20160659

14. Onji M, Sigl V, Lendl T, Novatchkova M, Ullate-Agote A, Andersson-Rolf A, et al. RANK drives structured intestinal epithelial expansion during pregnancy. Nature. 2025 Jan;637(8044):156–66. doi:10.1038/s41586-024-08284-1

15. Bando JK, Gilfillan S, Song C, McDonald KG, Huang SCC, Newberry RD, et al. The Tumor Necrosis Factor Superfamily Member RANKL Suppresses Effector Cytokine Production in Group 3 Innate Lymphoid Cells. Immunity. 2018 Jun 19;48(6):1208–1219.e4. doi:10.1016/j.immuni.2018.04.012

16. Xu H, Wang Y, Wang W, Fu YX, Qiu J, Shi Y, et al. ILC3s promote intestinal tuft cell hyperplasia and anthelmintic immunity through RANK signaling. Sci Immunol. 2025 May 16;10(107):eadn1491. doi:10.1126/sciimmunol.adn1491

17. Wallace KN, Pack M. Unique and conserved aspects of gut development in zebrafish. Dev Biol. 2003 Mar 1;255(1):12–29. doi:10.1016/s0012-1606(02)00034-9

18. Willms RJ, Foley E. Mechanisms of epithelial growth and development in the zebrafish intestine. Biochem Soc Trans. 2023 Jun 28;51(3):1213–24. doi:10.1042/BST20221375

19. Crosnier C, Vargesson N, Gschmeissner S, Ariza-McNaughton L, Morrison A, Lewis J. Delta-Notch signalling controls commitment to a secretory fate in the zebrafish intestine. Dev Camb Engl. 2005 Mar;132(5):1093–104. doi:10.1242/dev.01644

20. Wallace KN, Akhter S, Smith EM, Lorent K, Pack M. Intestinal growth and differentiation in zebrafish. Mech Dev. 2005 Feb;122(2):157–73. doi:10.1016/j.mod.2004.10.009

21. Rawls JF, Samuel BS, Gordon JI. Gnotobiotic zebrafish reveal evolutionarily conserved responses to the gut microbiota. Proc Natl Acad Sci. 2004 Mar 30;101(13):4596–601. doi:10.1073/pnas.0400706101

22. Lickwar CR, Camp JG, Weiser M, Cocchiaro JL, Kingsley DM, Furey TS, et al. Genomic dissection of conserved transcriptional regulation in intestinal epithelial cells. PLoS Biol. 2017 Aug;15(8):e2002054. doi:10.1371/journal.pbio.2002054

23. Willms RJ, Jones LO, Hocking JC, Foley E. A cell atlas of microbe-responsive processes in the zebrafish intestine. Cell Rep. 2022 Feb 1;38(5):110311. doi:10.1016/j.celrep.2022.110311

24. Jones LO, Willms RJ, Xu X, Graham RDV, Eklund M, Shin M, et al. Single-cell resolution of the adult zebrafish intestine under conventional conditions and in response to an acute Vibrio cholerae infection. Cell Rep. 2023 Nov 28;42(11):113407. doi:10.1016/j.celrep.2023.113407

25. Sato A, Miyoshi S. Fine structure of tuft cells of the main excretory duct epithelium in the rat submandibular gland. Anat Rec. 1997 Jul;248(3):325–31. doi:10.1002/(SICI)1097-0185(199707)248:3%3C325::AID-AR4%3E3.0.CO;2-O

26. Trier JS, Allan CH, Marcial MA, Madara JL. Structural features of the apical and tubulovesicular membranes of rodent small intestinal tuft cells. Anat Rec. 1987 Sep;219(1):69–77. doi:10.1002/ar.1092190112

27. Buissant des Amorie JR, Betjes MA, Bernink JH, Hageman JH, Geurts VE, Begthel H, et al. Intestinal tuft cell subtypes represent successive stages of maturation driven by crypt-villus signaling gradients. Nat Commun. 2025 Jul 22;16(1):6765. doi:10.1038/s41467-025-61878-9

28. Boyce BF, Xing L. Functions of RANKL/RANK/OPG in bone modeling and remodeling. Arch Biochem Biophys. 2008 May 15;473(2):139–46. doi:10.1016/j.abb.2008.03.018

29. ILC3s promote intestinal tuft cell hyperplasia and anthelmintic immunity through RANK signaling [Internet]. [cited 2025 Jul 29]. Available from: https://www.science.org/doi/10.1126/sciimmunol.adn1491 doi:10.1126/sciimmunol.adn1491

30. Feng X, Andersson T, Flüchter P, Gschwend J, Berest I, Muff JL, et al. Tuft cell IL-17RB restrains IL-25 bioavailability and reveals context-dependent ILC2 hypoproliferation. Nat Immunol. 2025 Apr;26(4):567–81. doi:10.1038/s41590-025-02104-y

31. Wang Z, Du J, Lam SH, Mathavan S, Matsudaira P, Gong Z. Morphological and molecular evidence for functional organization along the rostrocaudal axis of the adult zebrafish intestine. BMC Genomics. 2010 Jun 22;11:392. doi:10.1186/1471-2164-11-392

32. Malonga T, Vialaneix N, Beaumont M. BEST4+ cells in the intestinal epithelium. Am J Physiol-Cell Physiol. 2024 May;326(5):C1345–52. doi:10.1152/ajpcell.00042.2024

33. Wen J, Mercado GP, Volland A, Doden HL, Lickwar CR, Crooks T, et al. Fxr signaling and microbial metabolism of bile salts in the zebrafish intestine. Sci Adv. 2021 Jul 23;7(30):eabg1371. doi:10.1126/sciadv.abg1371

34. Sur A, Wang Y, Capar P, Margolin G, Prochaska MK, Farrell JA. Single-cell analysis of shared signatures and transcriptional diversity during zebrafish development. Dev Cell. 2023 Dec 18;58(24):3028–3047.e12. doi:10.1016/j.devcel.2023.11.001

35. Sur A, Segal EX, Nunneley MP, Sinclair JW, Prochaska MK, Dye LE, et al. Developmental regulation of intestinal best4+ cells. BioRxiv Prepr Serv Biol. 2025 Dec 19;2025.12.17.694935. doi:10.64898/2025.12.17.694935

36. Burclaff J, Bliton RJ, Breau KA, Ok MT, Gomez-Martinez I, Ranek JS, et al. A Proximal-to-Distal Survey of Healthy Adult Human Small Intestine and Colon Epithelium by Single-Cell Transcriptomics. Cell Mol Gastroenterol Hepatol. 2022;13(5):1554–89. doi:10.1016/j.jcmgh.2022.02.007

37. Elmentaite R, Kumasaka N, Roberts K, Fleming A, Dann E, King HW, et al. Cells of the human intestinal tract mapped across space and time. Nature. 2021 Sep;597(7875):250–5. doi:10.1038/s41586-021-03852-1

38. Zagoren E, Dias N, Santos AK, Smith ZD, Ameen NA, Sumigray K. Evidence of secondary Notch signaling within the rat small intestine. Dev Camb Engl. 2025 Jun 3;152(11):dev204277. doi:10.1242/dev.204277

39. Wang D, Spoelstra WK, Lin L, Akkerman N, Krueger D, Dayton T, et al. Interferon-responsive intestinal BEST4/CA7+ cells are targets of bacterial diarrheal toxins. Cell Stem Cell. 2025 Apr 3;32(4):598–612.e5. doi:10.1016/j.stem.2025.02.003

40. Mitchell KC, Withey JH. Danio rerio as a Native Host Model for Understanding Pathophysiology of Vibrio cholerae. In: Sikora AE, editor. Vibrio Cholerae: Methods and Protocols [Internet]. New York, NY: Springer; 2018 [cited 2025 Jul 3]. p. 97–102. Available from: 10.1007/978-1-4939-8685-9_9 doi:10.1007/978-1-4939-8685-9_9

41. Ito G, Okamoto R, Murano T, Shimizu H, Fujii S, Nakata T, et al. Lineage-specific expression of bestrophin-2 and bestrophin-4 in human intestinal epithelial cells. PloS One. 2013;8(11):e79693. doi:10.1371/journal.pone.0079693

42. Elmentaite R, Kumasaka N, Roberts K, Fleming A, Dann E, King HW, et al. Cells of the human intestinal tract mapped across space and time. Nature. 2021 Sep;597(7875):250–5. doi:10.1038/s41586-021-03852-1

43. Luna Velez MV, Neikes HK, Snabel RR, Quint Y, Qian C, Martens A, et al. ONECUT2 regulates RANKL-dependent enterocyte and microfold cell differentiation in the small intestine; a multi-omics study. Nucleic Acids Res. 2023 Feb 22;51(3):1277–96. doi:10.1093/nar/gkac1236

44. Hernández PP, Strzelecka PM, Athanasiadis EI, Hall D, Robalo AF, Collins CM, et al. Single-cell transcriptional analysis reveals ILC-like cells in zebrafish. Sci Immunol. 2018 Nov 16;3(29):eaau5265. doi:10.1126/sciimmunol.aau5265

45. Morales RA, Rabahi S, Diaz OE, Salloum Y, Kern BC, Westling M, et al. Interleukin-10 regulates goblet cell numbers through Notch signaling in the developing zebrafish intestine. Mucosal Immunol. 2022 Aug 1;15(5):940–51. doi:10.1038/s41385-022-00546-3

46. Bottiglione F, Dee CT, Lea R, Zeef LAH, Badrock AP, Wane M, et al. Zebrafish IL-4–like Cytokines and IL-10 Suppress Inflammation but Only IL-10 Is Essential for Gill Homeostasis. J Immunol Author Choice. 2020 Aug 15;205(4):994–1008. doi:10.4049/jimmunol.2000372

47. O’Leary CE, Ma Z, Culpepper T, Novak SW, DelGiorno KE. New insights into tuft cell formation: Implications for structure–function relationships. Curr Opin Cell Biol. 2022 Jun 1;76:102082. doi:10.1016/j.ceb.2022.102082

48. Angkasekwinai P, Srimanote P, Wang YH, Pootong A, Sakolvaree Y, Pattanapanyasat K, et al. Interleukin-25 (IL- 25) Promotes Efficient Protective Immunity against Trichinella spiralis Infection by Enhancing the Antigen- Specific IL-9 Response. Infect Immun. 2013 Oct;81(10):3731–41. doi:10.1128/IAI.00646-13.

49. Owyang AM, Zaph C, Wilson EH, Guild KJ, McClanahan T, Miller HRP, et al. Interleukin 25 regulates type 2 cytokine-dependent immunity and limits chronic inflammation in the gastrointestinal tract. J Exp Med. 2006 Apr 17;203(4):843–9. doi:10.1084/jem.20051496

50. Elmentaite R, Kumasaka N, Roberts K, Fleming A, Dann E, King HW, et al. Cells of the human intestinal tract mapped across space and time. Nature. 2021 Sep;597(7875):250–5. doi:10.1038/s41586-021-03852-1

51. Inaba A, Arinaga A, Tanaka K, Endo T, Hayatsu N, Okazaki Y, et al. Interleukin-4 Promotes Tuft Cell Differentiation and Acetylcholine Production in Intestinal Organoids of Non-Human Primate. Int J Mol Sci. 2021 Jul 24;22(15):7921. doi:10.3390/ijms22157921

52. Billipp TE, Fung C, Webeck LM, Sargent DB, Gologorsky MB, Chen Z, et al. Tuft cell-derived acetylcholine promotes epithelial chloride secretion and intestinal helminth clearance. Immunity. 2024 Jun 11;57(6):1243–1259.e8. doi:10.1016/j.immuni.2024.03.023

53. Ndjim M, Gasmi I, Herbert F, Joséphine C, Bas J, Lamrani A, et al. Tuft cell acetylcholine is released into the gut lumen to promote anti-helminth immunity. Immunity. 2024 Jun 11;57(6):1260–1273.e7. doi:10.1016/j.immuni.2024.04.018

54. Wrzosek L, Miquel S, Noordine ML, Bouet S, Chevalier-Curt MJ, Robert V, et al. Bacteroides thetaiotaomicron and Faecalibacterium prausnitziiinfluence the production of mucus glycans and the development of goblet cells in the colonic epithelium of a gnotobiotic model rodent. BMC Biol. 2013 May 21;11(1):61. doi:10.1186/1741-7007-11-61

55. Troll JV, Hamilton MK, Abel ML, Ganz J, Bates JM, Stephens WZ, et al. Microbiota promote secretory cell determination in the intestinal epithelium by modulating host Notch signaling. Development. 2018 Feb 23;145(4):dev155317. doi:10.1242/dev.155317

56. de Lau W, Kujala P, Schneeberger K, Middendorp S, Li VSW, Barker N, et al. Peyer’s patch M cells derived from Lgr5(+) stem cells require SpiB and are induced by RankL in cultured ‘miniguts’. Mol Cell Biol. 2012 Sep;32(18):3639–47. doi:10.1128/MCB.00434-12

57. Hickey JW, Becker WR, Nevins SA, Horning A, Perez AE, Zhu C, et al. Organization of the human intestine at single-cell resolution. Nature. 2023 Jul;619(7970):572–84. doi:10.1038/s41586-023-05915-x

58. Busslinger GA, Weusten BLA, Bogte A, Begthel H, Brosens LAA, Clevers H. Human gastrointestinal epithelia of the esophagus, stomach, and duodenum resolved at single-cell resolution. Cell Rep. 2021 Mar 9;34(10):108819. doi:10.1016/j.celrep.2021.108819

59. Wang G, Wang F, Meng Z, Wang N, Zhou C, Zhang J, et al. Uncovering potential genes in colorectal cancer based on integrated and DNA methylation analysis in the gene expression omnibus database. BMC Cancer. 2022 Feb 3;22(1):138. doi:10.1186/s12885-022-09185-0

60. Qu H, Su Y, Yu L, Zhao H, Xin C. Wild-type p53 regulates OTOP2 transcription through DNA loop alteration of the promoter in colorectal cancer. FEBS Open Bio. 2019 Jan;9(1):26–34. doi:10.1002/2211-5463.12554 PubMed PMID: 30652071;

61. He XS, Ye WL, Zhang YJ, Yang XQ, Liu F, Wang JR, et al. Oncogenic potential of BEST4 in colorectal cancer via activation of PI3K/Akt signaling. Oncogene. 2022 Feb;41(8):1166–77. doi:10.1038/s41388-021-02160-2

62. Li H, Wang X, Wang Y, Zhang M, Hong F, Wang H, et al. Cross-species single-cell transcriptomic analysis reveals divergence of cell composition and functions in mammalian ileum epithelium. Cell 2022 May 5;11(1):19. doi:10.1186/s13619-022-00118-7

63. Tang W, Zhong Y, Wei Y, Deng Z, Mao J, Liu J, et al. Ileum tissue single-cell mRNA sequencing elucidates the cellular architecture of pathophysiological changes associated with weaning in piglets. BMC Biol. 2022 May 30;20(1):123. doi:10.1186/s12915-022-01321-3

64. Westfall AK, Gopalan SS, Kay JC, Tippetts TS, Cervantes MB, Lackey K, et al. Single-cell resolution of intestinal regeneration in pythons without crypts illuminates conserved vertebrate regenerative mechanisms. Proc Natl Acad Sci. 2024 Oct 22;121(43):e2405463121. doi:10.1073/pnas.2405463121

65. Dougall WC, Glaccum M, Charrier K, Rohrbach K, Brasel K, De Smedt T, et al. RANK is essential for osteoclast and lymph node development. Genes Dev. 1999 Sep 15;13(18):2412–24. doi:10.1101/gad.13.18.2412

66. Akiyama T, Shimo Y, Yanai H, Qin J, Ohshima D, Maruyama Y, et al. The tumor necrosis factor family receptors RANK and CD40 cooperatively establish the thymic medullary microenvironment and self-tolerance. Immunity. 2008 Sep 19;29(3):423–37. doi:10.1016/j.immuni.2008.06.015

67. Kong YY, Yoshida H, Sarosi I, Tan HL, Timms E, Capparelli C, et al. OPGL is a key regulator of osteoclastogenesis, lymphocyte development and lymph-node organogenesis. Nature. 1999 Jan 28;397(6717):315–23. doi:10.1038/16852

68. Rossi SW, Kim MY, Leibbrandt A, Parnell SM, Jenkinson WE, Glanville SH, et al. RANK signals from CD4(+)3(-) inducer cells regulate development of Aire-expressing epithelial cells in the thymic medulla. J Exp Med. 2007 Jun 11;204(6):1267–72. doi:10.1084/jem.20062497

69. Hoshijima K, Jurynec MJ, Klatt Shaw D, Jacobi AM, Behlke MA, Grunwald DJ. Highly Efficient CRISPR-Cas9- Based Methods for Generating Deletion Mutations and F0 Embryos that Lack Gene Function in Zebrafish. Dev Cell. 2019 Dec 2;51(5):645–657.e4. doi:10.1016/j.devcel.2019.10.004

70. Schindelin J, Arganda-Carreras I, Frise E, Kaynig V, Longair M, Pietzsch T, et al. Fiji: an open-source platform for biological-image analysis. Nat Methods. 2012 Jun 28;9(7):676–82. doi:10.1038/nmeth.2019

71. Ewels PA, Peltzer A, Fillinger S, Patel H, Alneberg J, Wilm A, et al. The nf-core framework for community- curated bioinformatics pipelines. Nat Biotechnol. 2020 Mar;38(3):276–8. doi:10.1038/s41587-020-0439-x

72. Patel H, Manning J, Ewels P, Garcia MU, Peltzer A, Hammarén R, et al. nf-core/rnaseq: nf-core/rnaseq v3.19.0 - Tungsten Turtle [Internet]. Zenodo; 2025 [cited 2025 Aug 15]. Available from: https://zenodo.org/records/15631172 doi:10.5281/zenodo.15631172

73. Patro R, Duggal G, Love MI, Irizarry RA, Kingsford C. Salmon provides fast and bias-aware quantification of transcript expression. Nat Methods. 2017 Apr;14(4):417–9. doi:10.1038/nmeth.4197

74. Love MI, Huber W, Anders S. Moderated estimation of fold change and dispersion for RNA-seq data with DESeq2. Genome Biol. 2014 Dec 5;15(12):550. doi:10.1186/s13059-014-0550-8

75. Huang DW, Sherman BT, Lempicki RA. Systematic and integrative analysis of large gene lists using DAVID bioinformatics resources. Nat Protoc. 2009;4(1):44–57. doi:10.1038/nprot.2008.211

76. Sherman BT, Hao M, Qiu J, Jiao X, Baseler MW, Lane HC, et al. DAVID: a web server for functional enrichment analysis and functional annotation of gene lists (2021 update). Nucleic Acids Res. 2022 Jul 5;50(W1):W216–21. doi:10.1093/nar/gkac194

77. Hao Y, Stuart T, Kowalski MH, Choudhary S, Hoffman P, Hartman A, et al. Dictionary learning for integrative, multimodal and scalable single-cell analysis. Nat Biotechnol. 2024 Feb;42(2):293–304. doi:10.1038/s41587-023-01767-y

